# Redefining the architecture of ferlin proteins: insights into multi-domain protein structure and function

**DOI:** 10.1101/2022.01.18.476802

**Authors:** Matthew J. Dominguez, Jon J. McCord, R. Bryan Sutton

## Abstract

Ferlins are complex, multidomain proteins whose functions involve membrane fusion, membrane repair, and exocytosis. Progress in studying the domain composition of ferlins has been problematic due to the large size of the ferlin proteins and the difficulty in computing accurate residue ranges for each respective ferlin protein domain. However, recent advances in *in silico* protein folding methods have significantly enhanced the understanding of the domain structure of these complex proteins. To compute unbiased domain boundaries, we used RoseTTAFold to assemble full-length models for each of the six human ferlin proteins (dysferlin, myoferlin, otoferlin, Fer1L4, Fer1L5, and Fer1L6). Despite the differences in amino acid sequence between the ferlin proteins, the domain ranges and the distinct subdomains in some of the ferlin domains were remarkably consistent. Further, the RoseTTAFold/AlphaFold2 *in silico* boundary prediction methods allowed us to describe and characterize a previously unknown C2 domain, ubiquitous in all human ferlins, which we refer to as C2-FerA. At present, the ferlin domain-domain interactions implied by the full-length *in silico* models are predicted to have a low accuracy; however, the use of RoseTTAFold and AlphaFold2 as a domain finder has proven to be a powerful research tool in understanding ferlin structure.

## Introduction

Ferlins are large, multi-domain proteins that mediate Ca^2+^-dependent phospholipid interactions in multiple cell types [1]. In humans six paralogous ferlin genes have been annotated within the genome: dysferlin (Fer1L1), otoferlin (Fer1L2), myoferlin (Fer1L3), Fer1L4, Fer1L5, and Fer1L6 [1]. Three of the ferlin proteins, dysferlin, otoferlin, and myoferlin, have well-studied biological functions. Dysferlin is an important component in skeletal muscle membrane repair, and mutations have been implicated in various muscular dystrophies [2–4]. Otoferlin is expressed in the brain and in vestibular hair cells in the ear [5], and is involved in Ca^2+^-dependent exocytosis in hair cells [6, 7] Mutations in the otoferlin gene have been linked to non-syndromic recessive sensorineural hearing loss in humans [8]. Myoferlin is required for normal muscle cell development [9] and has established links to metastatic cancers [10, 11]. Fer1L4 has been categorized as a pseudogene in humans, and it may only function as a long noncoding RNA (lncRNA) [12–18]; however, Fer1L4 may act as a functional protein in mice and zebrafish [1]. There is some evidence that Fer1L5 [19–21] and Fer1L6 [22–24] are active proteins in humans, but the exact function of these ferlins is currently unknown.

The defining feature of ferlin proteins is their multiple tandem C2 domains. The prototypical C2 domain is approximately 130 residues in length, possesses eight *β*-strands that fold around a central Greek key motif with two *β*-sheets that form a *β*-sandwich [25]. There are at least two known folding topologies for C2 domains, Type-I (Type-S) and Type-II (Type-P) [26]. Three loops at the apex of the domain form a cup-like Ca^2+^ binding pocket, and are labeled loops 1, 2, and 3 [27]. While not all C2 domains bind Ca^2+^, the C2 domains that do bind Ca^2+^, such as the two tandem C2 domains of synaptotagmin-1, possess up to six conserved acidic residues within the binding pocket with which to coordinate divalent cations [28]. Hydrophobic residues at the apices of loops 1 and 3 of the Ca^2+^ binding pocket form the basis for the Ca^2+^-dependent peripheral phospholipid interaction that is characteristic of many C2 domains [29].

Early analysis of dysferlin’s primary sequence concluded that there were at least two C2 domains in the protein [30]. Later analyses extended the complexity of the ferlins to include six to seven C2 domains of mixed topology [31, 32]. After these initial estimates, several studies published differing domain boundaries and topologies for the C2 domains of dysferlin [33–35]. The assumptions used to establish the early dysferlin domain definitions were based on the known fold of the classical C2 domains such as synaptotagmin-1 C2A and C2B; however, primary and secondary sequence alignments used to determine domain boundaries could not precisely rationalize synaptotagmin-like C2 domain boundaries in all ferlin C2 domains. The synaptotagmin-like C2 domain assumption for ferlin proteins led to domain predictions with an incorrect number of *β*-strands per C2 domains and a ferlin molecule dominated by unstructured linker sequences. With the advent of AlphaFold2 and RoseTTAFold [36, 37], we can now accurately compute unbiased C2 domain boundaries in large complex proteins such as ferlins (**Fig. 1**). Furthermore, previously unknown domains can be defined within these proteins. While the domain-domain interactions modeled for the various ferlins using AlphaFold2 and RoseTTAFold are likely not yet accurate, these two complementary methods make excellent domain finders that can produce experimentally testable domain boundaries to study complex multi-domain protein such as ferlins.

**Fig 1.**
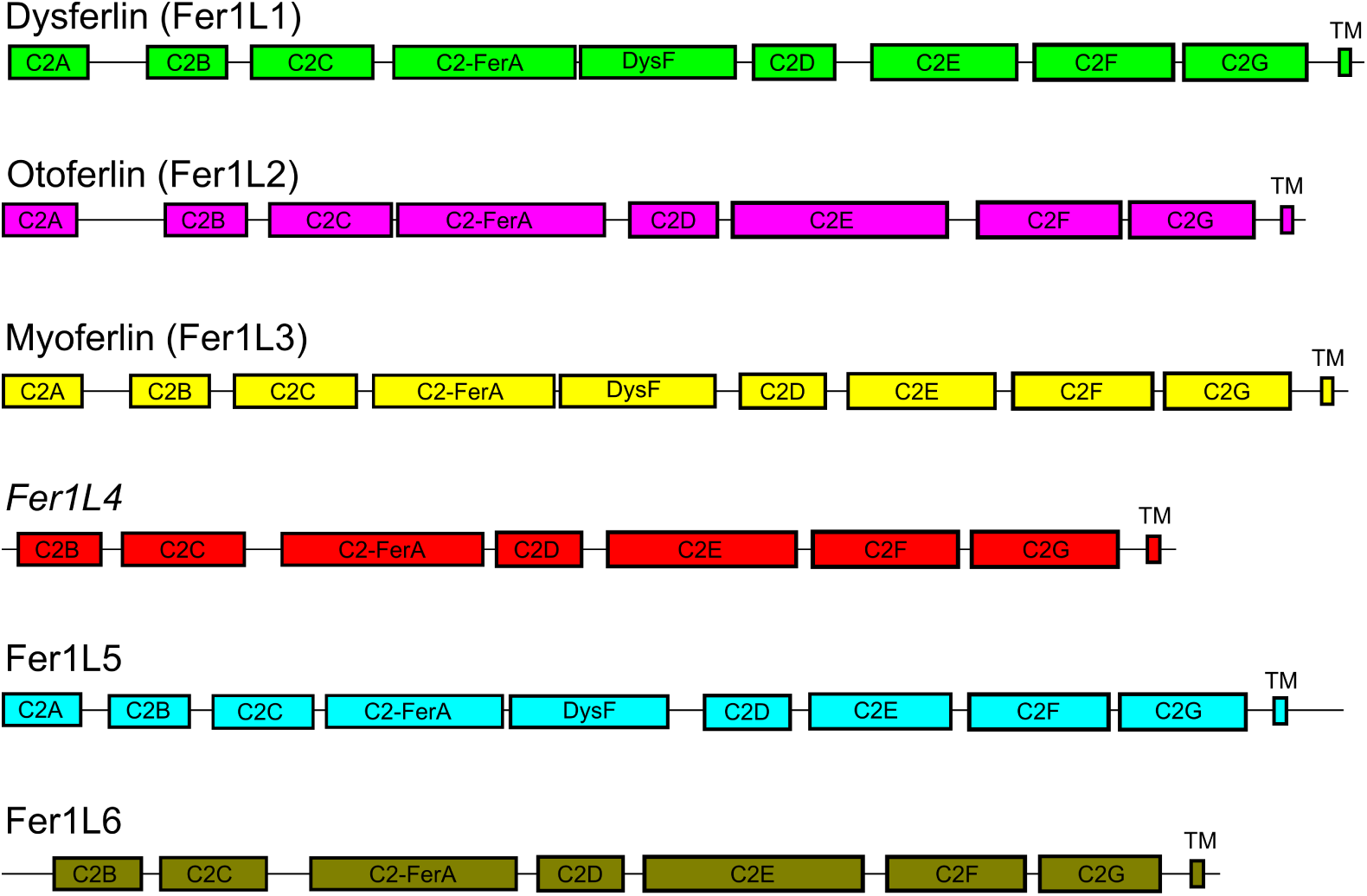
Scaled schematic diagram of the human ferlin family members. The length of the total protein is in proportion to the length of each domain. \**Fer1L4* in humans is currently characterized as a pseudogene. ‘TM’ indicates the transmembrane span.

## Materials and methods

### Model Building and Domain Definitions

Overlapping primary sequences of each of the ferlin family members were submitted to the Robetta server for structural assessment using the RoseTTAFold algorithm. Assuming that there is little to no cooperative folding between domains, the amino acid sequence of each domain would be expected to fold independently [38, 39], thereby providing an internal control for the folding results. Overlapping domains were superimposed, and a model of the entire molecule was assembled using Modeller [40]. We utilized protein sequence from the UniProt database (dysferlin, O75923; myoferlin, Q9NZM1; otoferlin, Q9HC10; Fer1L4, A9Z1Z3; Fer1L5, A0AVI2; and Fer1L6, Q2WGJ9). The final assembled protein models were relaxed using Pyrosetta to optimize side chain interactions in preparation for molecular dynamics. The energy minimization script used is reproduced in the supplementary section (**Fig. 1**).

C2 domains are defined as protein domains possessing eight *β*-strands that fold with a characteristic C2 domain topology [41]. The boundary condition for each C2 domain in each modeled ferlin protein is defined as a range of residues that begin with the first residue that contributes to the backbone hydrogen bond pattern of the first *β*-strand of the domain through to the last residue that contributes a backbone hydrogen bond to the eighth *β*-strand of the domain. Loops that significantly deviate from the typical Type-II C2 pattern, as defined by the C2A domain of dysferlin, and are unique to the ferlin family were defined as subdomains. A more extensive definition of each subdomain is provided in the results section.

### Molecular Dynamics

Molecular dynamics (MD) systems were prepared using Charmm-GUI [42] to regularize the geometry of the extracted C2 domain models from the full-length ferlin molecules. Individual domains were simulated in an explicit water box using NAMD [43]. The MD was run for 50 ns to relieve clashes and to improve any nonideal stereochemistry that resulted from the model building process. The CHARMM36m force field was used throughout. Simulations of the DysF regions of dysferlin, myoferlin, and Fer1L5 were performed using WYF parameters for pi-cation interactions. The protein systems were solvated in a cubic water box (having basis vector lengths of ∼80 Å) under periodic boundary conditions using the TIP3 water model. The total charge of the system was neutralized by randomly substituting water molecules with K^+^ and Cl^-^, in addition to a 0.15 M overall salt concentration. The initial energy minimization was performed for 10,000 steps to avoid any bad contacts generated while solvating the system. A switching function was applied to the van der Waal’s potential energy from 10 to 12 Å to calculate nonbonded interactions. The Particle Mesh Ewald (PME) algorithm was used to calculate electrostatic interactions. Equilibration runs used the NVT ensemble at 300 K. Molecular dynamics production runs were performed under NPT conditions at 303.15 K. The domain models were extracted and analyzed using VMD.

### Model Assessment

Ten aligned snapshots were extracted from the MD simulations. From the set of 10 snapshots, a representative model of each domain was selected based on its near-zero QMEAN Z-score as computed by SWISS-MODEL [44]. Violations in Ramachandran angles, clash scores, and the MolProbity scores for each predicted domain are reported in Supplementary Data (S1, S2, S3, S4, S5, S6).

### Model-based Primary Sequence Analysis

The structures and models of the known C2 domains of ferlins were superimposed and aligned using PROMALS3D [45]. The primary sequence alignment from PROMALS3D was used to compute the mean similarity and identity fractions using the SIAS server (http://imed.med.ucm.es/Tools/sias.html) for each domain Supplementary Data (**Fig. S1, S2, S3, S4, S5, S6, S7, S8, S9**). Alignment graphics in the Supplementary Material were prepared using ESPript [46].

### Cloning, Expression, Purification, and Characterization of C2-FerA

The gene with codon-optimized human dysferlin cDNA was obtained from Addgene (Plasmid 67878) [47]. The DNA sequence corresponding to the human dysferlin C2-FerA domain (residues 588 - 868) was subcloned using In-Fusion® Snap Assembly kit into a pET28a expression vector with an N-terminal 6 x His-tagged maltose-binding protein (MBP) fusion tag and a tobacco etch virus (TEV) cleavage site. The human dysferlin C2-FerA domain expression plasmid was transformed into Mix & Go BL21 Gold (DE3) competent cells and plated on LB-agar plates supplemented with 50 *µ*g/mL kanamycin and incubated overnight at 37 °CMultiple colonies were used to inoculate 100 mL of sterile Luria Broth (LB) containing 50 *µ*g/mL kanamycin and were left in a shaking incubator overnight at 250 rpm and 37 °C for 12-16 hours. 15 mL of the LB starter culture was used to inoculate a 2800 mL baffled Fernbach flask containing 1 L of sterile terrific broth (TB) supplemented with 50 *µ*g/mL kanamycin and was left in a shaking incubator at 250 rpm and 37 °C for four hours or until the culture reached an OD_600_ of at least 2.0. Next, the shaking incubator was cooled to 18 °C, and protein expression was induced by the addition of 400 *µ*L of 1 M IPTG. After induction, the culture was left in the shaking incubator (250 rpm and 18 °C) for 24 hours. The culture was harvested by centrifugation and the cell pellets were flash-frozen in liquid nitrogen and stored at -80 °C until needed.

To begin the purification, cells were thawed in lysis buffer (40 mM HEPES pH 7.4, 150 mM NaCl), ruptured using a microfluidizer, and spun using a Beckman JA-20 rotor at 19,500 rpm and 4 °C for 1 hour. The supernatant was applied to a Ni-IDA (His60 Ni Superflow Resin) affinity column, which was equilibrated with lysis buffer, and allowed to bind overnight at 4 °C . The supernatant was allowed to pass through the column and 200 mL lysis buffer including 20 mM imidazole was used to wash the column. Finally, human dysferlin C2-FerA was eluted with 35 mL lysis buffer including 250 mM imidazole. The resulting fusion protein was cleaved with 4 mg of Tobacco Etch Virus Protease (TEV) overnight at 4 °C . Next, the TEV cleaved protein was loaded onto a Q Sepharose® column using a Biorad NCG. Most of the MBP protein did not bind to the Q Sepharose® column. C2-FerA was eluted from the ion exchange resin by applying an NaCl gradient. The fractions containing the C2-FerA domain were pooled and concentrated for size exclusion chromatography using Superdex-75 resin. Purity was assessed using SDS PAGE Any-kD Mini-PROTEAN TGX Stain-Free gels from Biorad and protein concentration was quantified using absorbance at 280 nm and the calculated extinction coefficient (**Fig. 6A**).

Purified C2-FerA was buffer exchanged into a 10 mM sodium phosphate pH 7.4 buffer for circular dichroism (CD) spectroscopy using a J-850 spectropolarimeter from JASCO Corp. The CD spectrum was collected from 250 - 180 nm with 4 *µ*M C2-FerA. Data collection was performed at room temperature with an acquisition rate of 1 nm/sec and a data pitch of 0.1 nm. Additionally, CD spectra of 10 mM sodium phosphate pH 7.4 alone were collected and subtracted from the C2-FerA CD spectra to produce the final CD spectra used for fitting. The C2-FerA buffer-subtracted CD spectra were normalized to mean residue ellipticity and the secondary structure was estimated using the BeStSel webserver [48, 49]. We used the PDBMD2CD webserver [50] to calculate CD spectra based on the human dysferlin C2-FerA model and to obtain an estimate of secondary structure. The calculated values obtained from PDBMD2CD were then compared with the experimental CD spectra and secondary structure estimates (**Fig. 6B**).

## Results

### Overall ferlin domain structure

An early prediction of the C2 domain topology in dysferlin suggested the presence of both Type-I and Type-II C2 domains [32]. The prediction of mixed C2 domain topologies likely had to do with the difficulty in fitting most of the domains into known C2 domain sequences or secondary structure patterns. Our ferlin models, using both RoseTTAFold and AlphaFold2, predict that all C2 domains of ferlin proteins are Type-II. Interestingly, our ferlin models also show long conserved insertions within the predicted loops of many of the C2 domains. We refer to these loop insertions as subdomains. Here, we define subdomains as any C2 domain loop insertion greater than 20 amino acids (relative to the reference dysferlin C2A domain) that possesses conserved secondary structural elements across at least two human ferlin paralogs.

### Analysis of ferlin C2A domains

Of the five known human ferlin proteins, dysferlin, otoferlin, myoferlin, and Fer1L5 possess domains identified as C2A [51] (**Fig. S1**). Fer1L4 and Fer1L6 begin with C2B domains [51] (**Fig. 2**). Most of the structural and functional data on ferlin domains has focused on the C2A domain because the boundaries were simpler to predict [52–55]. The first residue is the N-terminal residue of C2A, while the C-terminal C2A domain boundary can be predicted from its overall similarity to other C2 structures. Currently, there are high-resolution crystal structures and solution-state NMR spectroscopy structures for dysferlin C2A [52, 54], myoferlin C2A [53, 56], and otoferlin C2A [55]. Additionally, dysferlin [52, 54] and myoferlin C2A [53] have been solved with liganded divalent cations. Dysferlin (4IHB) and myoferlin (6EEL) C2A domains are similar in overall structure with a 1.91 Å RMSD between the two structures [53]. The C2A domain of otoferlin lacks loop 1 of its putative Ca^2+^-binding pocket, so it cannot coordinate Ca^2+^ [55]. Fer1L5 C2A is the only ferlin without experimental structural analysis; however, our current model of Fer1L5 can provide some predictive power to infer its properties (**Fig. 2**). Fer1L5 only has one acidic residue (Asp-69) out of six acidic residues in the Ca^2+^ binding region (CBR) typically used to coordinate Ca^2+^. Therefore, we predict that the human Fer1L5 C2A domain does not possess the capacity to bind Ca^2+^. *β*-strand 3 of the Fer1L5 C2A model has a poly-basic region similar to *β*-strand 3 of dysferlin C2A; therefore, it may possess the ability to bind to negatively charged phospholipid surfaces [57]). Loop 1 of Fer1L5 may bind phospholipid membranes via its hydrophobic residues, and loop 3 has several basic residues that may direct the coordination of negatively charged phospholipids.

**Fig 2.**
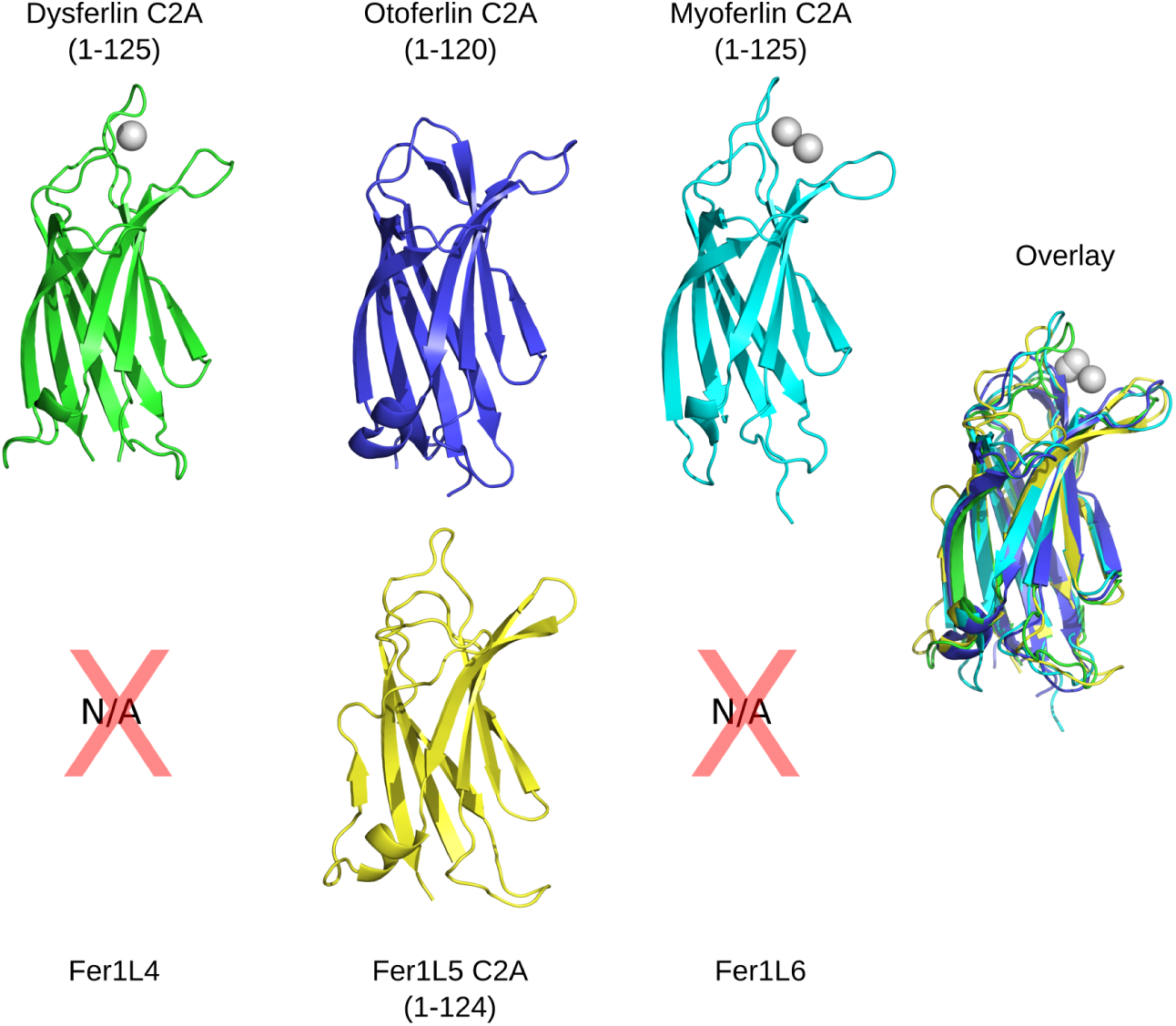
C2A structures and model of C2A from Fer1L5. Dysferlin C2A (green, 4IHB), otoferlin C2A (blue, 3L9B), myoferlin C2A (cyan, 6EEL), and Fer1L5 C2A (yellow). The gray spheres are the divalent cations from the crystal structures of dysferlin C2A (4IHB, chain E) and myoferlin C2A (6EEL). The amino acid boundaries from each respective C2 domain are listed with the domain assignment. The superposition of all four C2A domains are labeled as ‘Overlay’. The ferlins without a C2A domain are listed as Not Applicable (N/A).

### Analysis of ferlin C2A-C2B linkers

The linker between the C2A and C2B domains (A-B linker) is the longest linker sequence modeled in the human ferlin molecules. Our models predict this region to be mostly unstructured due to a high fraction of proline and glycine residues. For example, the A-B linker sequence in dysferlin is 93 amino acids in length and is 22% proline. The A-B linker sequence from myoferlin is 73 amino acids in length and is 13.7% proline and 13.7% glycine. Otoferlin has the longest A-B linker sequence in the ferlin family with 132 amino acids but possesses a lower fraction of proline and glycine, 8.3% and 9.1% respectively. Fer1L5 has the shortest A-B linker of the human ferlin family, 42 amino acids, and a lower fraction of proline and glycine, 4.8% and 7.1%, respectively. Despite the prevalence of proline and glycine in the A-B linkers among all ferlins, our RoseTTAFold models predict two *α*-helices near the center of the chain. The two helices are oppositely charged, and they are modeled as an electrostatically-coupled pair. The N-terminal helix tends to be negatively charged, while the C-terminal helix in the A-B linker tends to be positively charged. The A-B linker in dysferlin possesses two pathological mutations; P132L and A170E. P132L occurs within a proline-rich region of the linker near the C-terminus of the C2A domain. The A170E mutation occurs within the negatively charged helical region of the linker.

In dysferlin, there are 14 consensus phosphorylation sites predicted within the helices of the A-B linker [58]. Interestingly, the exon 5a insertion of dysferlin adds 31 residues to its A-B linker [59], and another nine potential phosphorylation sites. The prevalence of potential phosphorylation sites in the dysferlin A-B linker suggests that charge modification via serine/threonine phosphorylation may be a regulatory mechanism.

### Analysis of ferlin C2B domains

The reported boundaries for the C2B domain of dysferlin are the most consistent among all of the previous C2B domain predictions [1, 32, 34, 35] (**Fig. S2**). The six ferlin C2B models from RoseTTAFold superimpose on each other with an RMSD less than 1.0 Å (**Fig. 3**), suggesting a consistent fold between ferlin C2B domains. All human ferlin C2B domains closely resemble a prototypical C2 domain in both length and structure. The C2B domains of the ferlins generally lack the typical C2 domain Ca^2+^ binding motif; however, they possess putative lipid-binding residues on loops 1 and 3. Interestingly, the Type-2 ferlins (otoferlin, Fer1L4, and Fer1L6) possess a conserved histidine residue at the locus typically used for axial Ca^2+^ coordination in other divalent cation binding C2 domains.

**Fig 3.**
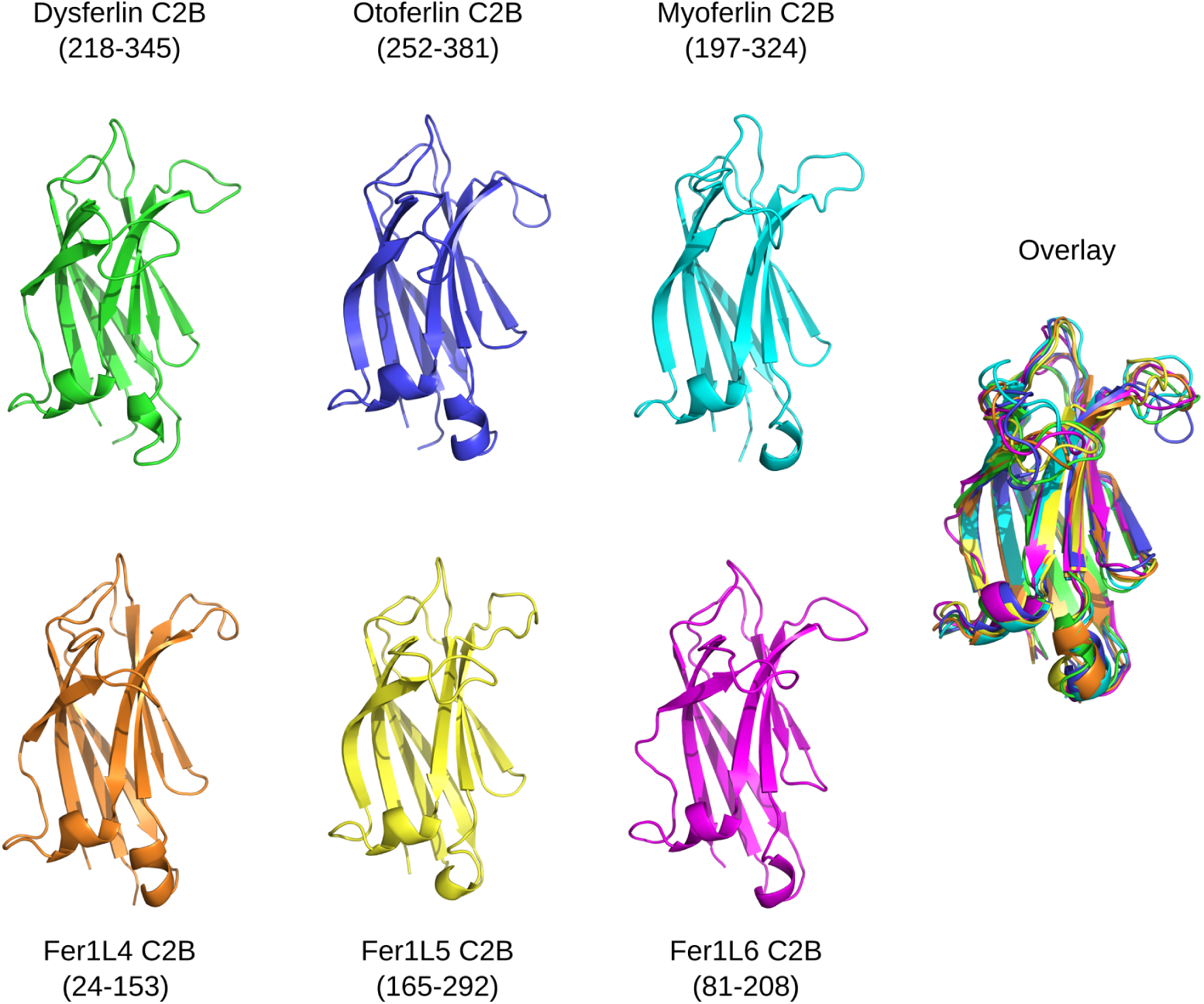
C2B models from dysferlin C2B (green), otoferlin C2B (blue), myoferlin C2B (cyan), Fer1L4 C2B (orange), Fer1L5 C2B (yellow), and Fer1L6 (magenta). Amino acid ranges for each domain are listed under the domain label. The superposition of all six C2B domains are labeled as ‘Overlay’.

### Analysis of ferlin C2C domains

The C2C domain is one of the most intricate ferlin C2 domains, since some ferlin C2C domains possess up to three subdomains within its boundaries (**Fig. 4**). The first subdomain of C2C occurs between *β*-strands 1 and 2 (*β*1-2), introducing a long, amphipathic *α*-helix within loop 1 of the C2 domain (**Fig. S3**), *α*1). The *β*1-2 subdomain of C2C is the only example of a loop 1 or *β*1-2 subdomain in the ferlin family. A loop 1 insertion is particularly notable as this loop typically participates in Ca^2+^-dependent phospholipid binding in many C2 domains. In fact, the *β*1-2 subdomain of C2C shares similarity to the loop 1 (CBR1) insertion observed in cytosolic phospholipase A2 [60]. We predict that C2C would possess a similar Ca^2+^ and phospholipid-binding activity as cytosolic phospholipase A2. In the Ca^2+^-dependent C2 domains, the domain itself contributes at least five of six total coordinating oxygen atoms to bind Ca^2+^; a phospholipid headgroup contributes the remaining coordinating oxygen. All six human ferlin C2C domains possess most of the requisite divalent cation coordination residues to bind Ca^2+^. Although the precise identification of the residues that could bind Ca^2+^ from the domain is not clear from modeling, the *β*1-2 subdomain could contribute acidic residues to complete the hexadentate Ca^2+^ coordination. Interestingly, five of the six ferlin C2C models are predicted to include the loop 1 subdomain; only Fer1L5 lacks this *β*1-2 subdomain.

**Fig 4.**
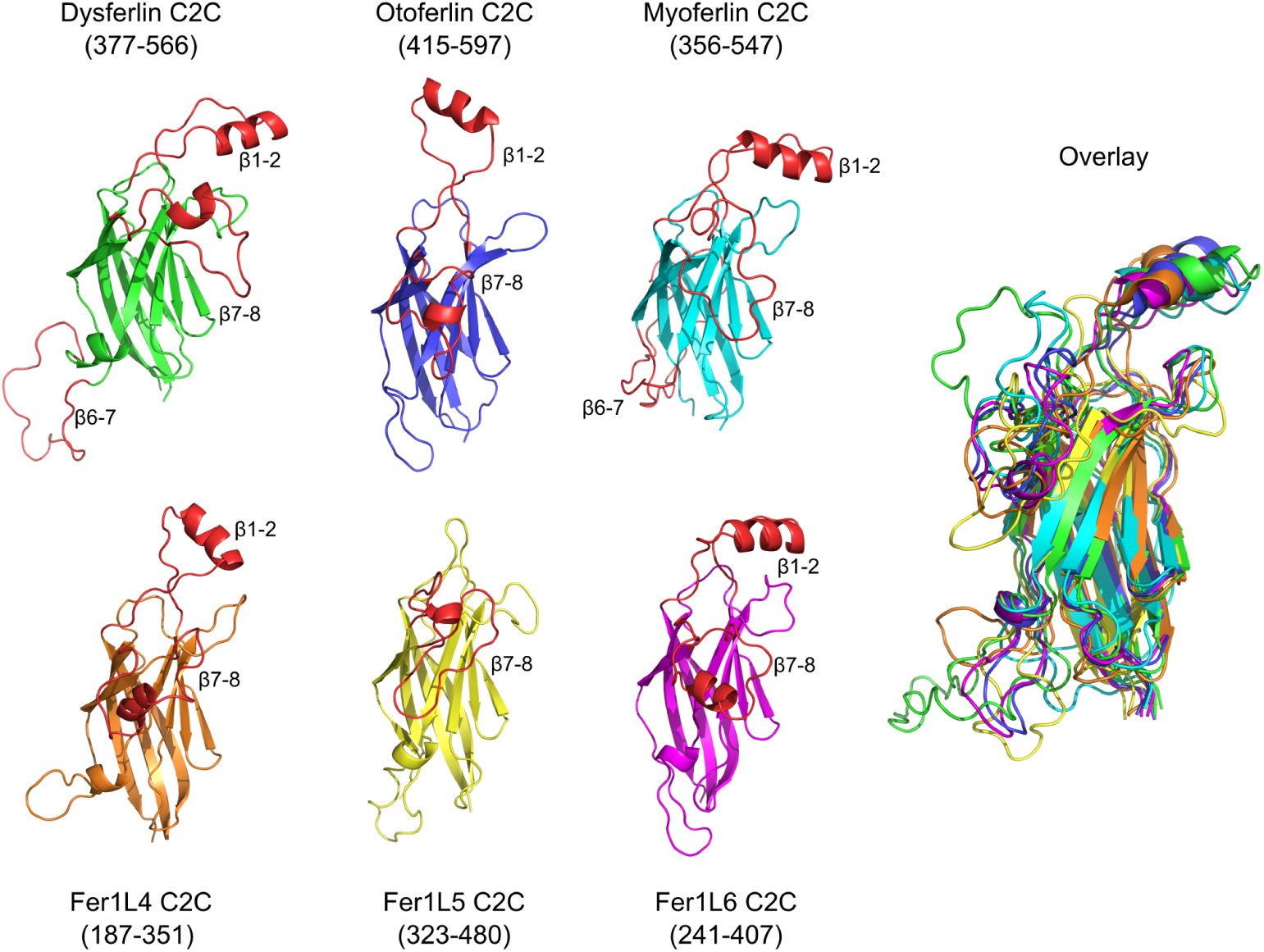
C2C models for dysferlin C2C (green), otoferlin C2C (blue), myoferlin C2C (cyan), Fer1L4 C2C (orange), Fer1L5 C2C(yellow), and Fer1L6 C2C (magenta). Amino acid ranges for each domain are listed under the domain label. The *β*1-2, *β*6-7, and *β*7-8 subdomains for C2C are labeled and shown in red. The superposition of all six C2C domains are labeled as ‘Overlay’.

**Fig 5.**
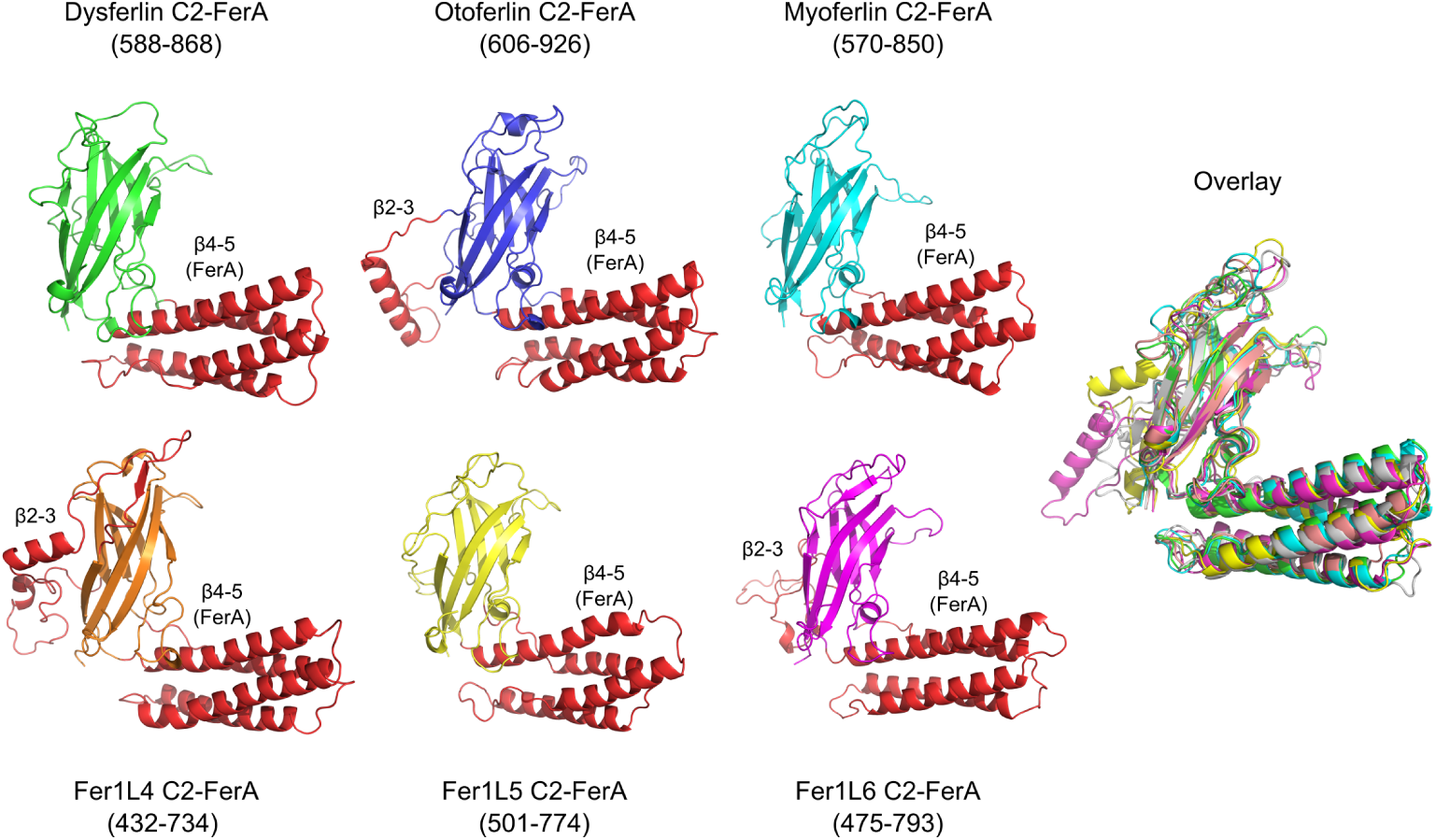
C2-FerA models for dysferlin C2-FerA (green), otoferlin C2-FerA (blue), myoferlin C2-FerA (cyan), Fer1L3 C2-FerA (orange), Fer1L5 C2-FerA (yellow), Fer1L6 C2-FerA (magenta). The FerA subdomain (*β*4-5) is shown as red *α*-helices, as is the *β*2-3 subdomain unique to Type-2 ferlins. The superposition of all six C2-FerA models are labeled as ‘Overlay’.

The C2C domains of dysferlin and myoferlin possess a unique, 30-amino acid long subdomain between *β*-strands 6-7 (*β*6-7). The overall structure of the *β*6-7 subdomain of C2C is uncertain; but, it may contain at least two *α*-helices. The exon 17 exclusion of dysferlin abbreviates the *β*6-7 subdomain to a single Val residue, making it a similar size to the other human ferlin C2C domains (**Fig. S3**) [59]. Interestingly, the exon 17 exclusion isoform of dysferlin seems to be characteristic of the dysferlin transcripts present in the blood; yet it is effectively absent in muscle tissue [59]. The third subdomain of C2C occurs between *β*-strands 7-8 (*β*7-8), and it is approximately 20 residues in length across all human ferlins. There is no obvious sequence similarity or discernible function that can be inferred from our models or alignment of *β*7-8 (**Fig. S3**, *α*3).

### Analysis of ferlin C2-FerA domains

The FerA subdomain was initially discovered based on the predicted *α*-helical propensity of the dysferlin amino acid region located between C2C and inner DysF [61]. The isolated FerA subdomain has been shown to interact with phospholipids in a Ca^2+^-dependent manner [61]; however, the exact mechanism and biological function of FerA is still under study. Prior to the availability of RoseTTAFold and AlphaFold2, we observed tell-tale *β*-strands from secondary structure analysis on either side of the FerA four-helix bundle [45]. Using the predicted flanking *β*-strands, we could assemble a complete C2 domain with the FerA four-helix bundle modeled as a subdomain between *β*-strand 4 and *β*-strand 5 (*β*4-5). The new C2 domain is also independently modeled in our AlphaFold2 and RoseTTAFold ferlin analysis, confirming our initial prediction (**Fig. 6**). We refer to this novel C2 domain as C2-FerA. C2-FerA was likely not discovered in the initial automated domain searches of the ferlins because the large four-helix bundle that interrupts the typical C2-like motif is nearly as large as a prototypical C2 domain (**Fig. S4**). C2-FerA is the largest domain by amino acid sequence in 5 out of the 6 human ferlins (**Fig. 1**). Interestingly, the Type-2 ferlins include an additional subdomain in C2-FerA that occurs between *β*-strands 2-3 (*β*2-3). The *β*2-3 subdomain is uniformly negatively charged in all three Type-2 ferlins.

**Fig 6.**
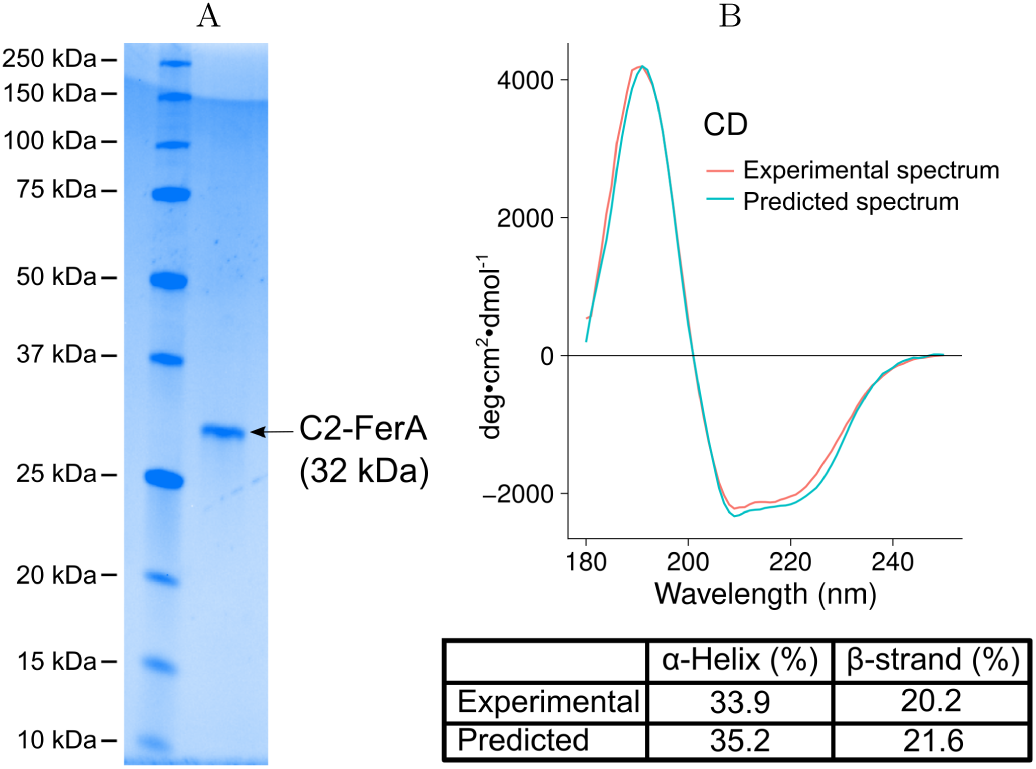
Dysferlin C2-FerA analysis. **A.** SDS-PAGE showing the purified dysferlin C2-FerA domain versus molecular weight size markers. **B.** Far-UV CD spectrum of purified dysferlin C2-FerA (red curve) and predicted C2-FerA spectrum derived from the model (blue curve). The inset table reports the secondary structure summary of the purified dysferlin C2-FerA domain (Experimental), and the dysferlin C2-FerA model (Predicted).

To validate our novel C2-FerA model, we cloned, expressed, and purified the human dysferlin C2-FerA domain based on our RoseTTAFold results (**Fig. 6A**). The secondary structure of the purified domain was measured by far-UV circular dichroism (CD) spectroscopy [49]. The experimental CD spectrum was consistent with a domain possessing a mix of *α*-helix and *β*-sheet secondary structural elements (**Fig. 6B**). We also predicted the CD spectra of our C2-FerA model and computed the contribution of its secondary structural elements [50]. Both the experimentally derived CD spectra and the computed CD spectra agreed exceptionally well (**Fig. 6B**); thus, confirming our C2-FerA domain model and validating the previously undescribed C2-FerA domain.

### Analysis of ferlin DysF regions

The DysF region is unique to the Type-1 ferlins, and is composed of two DysF domains located near the center of the ferlin protein’s primary sequence [51]. The structures of the inner DysF domains of dysferlin and myoferlin were solved by X-ray crystallography and solution-state NMR spectroscopy, respectively [62, 63]. From primary sequence alignments, the inner DysF domain appears to be embedded within the sequence of the outer DysF domain (**Fig. S5**) [51]; however, it was unclear how to resolve the folding discontinuity using only the primary sequence. Our molecular modeling provides a possible solution to the complex topology of the of the DysF region (**Fig. 7**). Indeed, there are two predicted DysF domains, one domain embedded within the other. The inner DysF domain of dysferlin and myoferlin is made up of two, contiguous anti-parallel *β*-strands, dominated by Trp-Arg *π*-cation interactions [62, 63]. The first *β*-strand of the outer DysF domain is located on the N-terminal side of the DysF region, while its second *β*-strand is located at the C-terminal end of the DysF region. The continuous inner DysF domain is embedded between the two halves of the outer DysF domain. The net result is two tandem DysF domains spaced apart by a single *α*-helix (**Fig. 7**). The helix between the two DysF domains in dysferlin, myoferlin, and Fer1L5 are all about 18-residues in length and possess no outstanding biochemical characteristics, suggesting that it may serve as a spacer between DysF domains.

**Fig 7.**
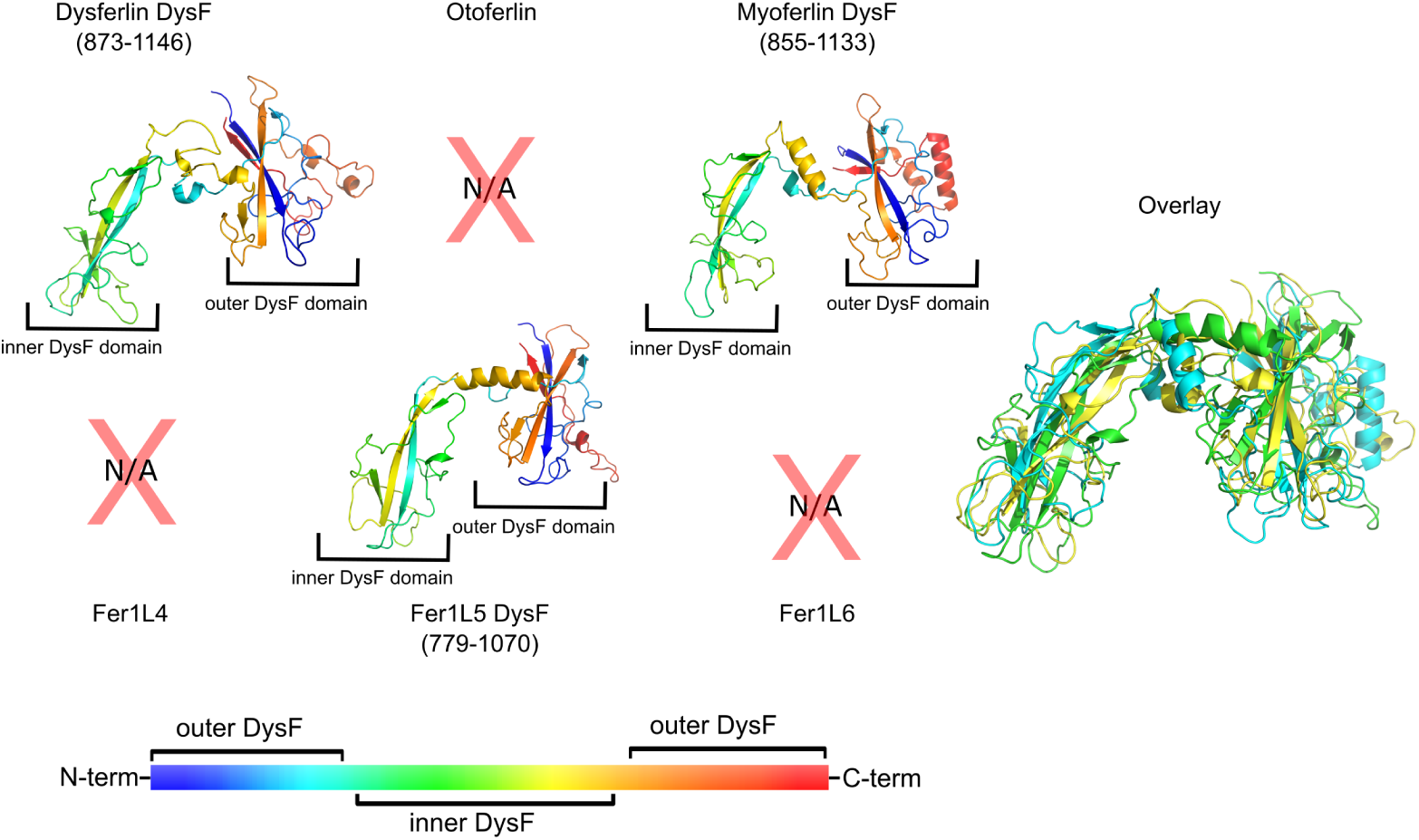
Models of the complete DysF region of the Type-1 ferlins; dysferlin, myoferlin, and Fer1L5. The inner DysF domain is positioned at the left of each model, whereas the outer domain is positioned on the right side. The models of the complete DysF region have been colored as a gradient from blue to red to highlight the crossover topology of the outer DysF domain’s fold. The color bar illustrates the progression of colors from N-terminus to C-terminus.

The physiological function of the DysF region of the Type-1 ferlins is unclear. There are examples of homologous DysF domains in *Saccharomyces cerevisiae* peroxisomal proteins Pex30p, Pex31p, and Pex32p where they are implicated in maintaining the number and size of peroxisomes [64, 65]. Additionally, the sporulation-specific protein 73 (SPO73) of *S. cerevisiae* is composed solely of a single DysF domain and is required for prospore membrane extension [66]. Interestingly, the DysF domain is only noted in one other human protein besides the Type-1 ferlins. The tectonin beta-propeller repeat-containing protein 1 (TECPR1) contains two DysF domains; however, the DysF domains are composed of continuous sequences and are not embedded as they are in the Type-1 ferlins. TECPR1 is involved in the fusion between autophagosomes and lysosomes, and it promotes the degradation of protein aggregates. Therefore, the DysF domain likely contributes to the membrane fusion/membrane binding activity of TECPR1 [67] and to the Type-1 ferlin proteins [68].

### Analysis of ferlin C2D domains

C2D resembles a prototypical C2 domain with no discernible subdomains (**Fig. 8**). Yet despite the straightforward 3D fold of C2D, there has been a wide variety of boundary estimates published for C2D [1, 32, 34, 35]. Five of the six C2D domains in the human ferlins possess all requisite residues typical of C2 domain Ca^2+^ coordination (**Fig. S6**). Indeed, the MIB (Metal Ion-Binding Site Prediction and Docking Server) server [69] predicts that Ca^2+^ can bind within the canonical divalent cation binding pocket of the ferlin C2D domains, expect Fer1L5.

**Fig 8.**
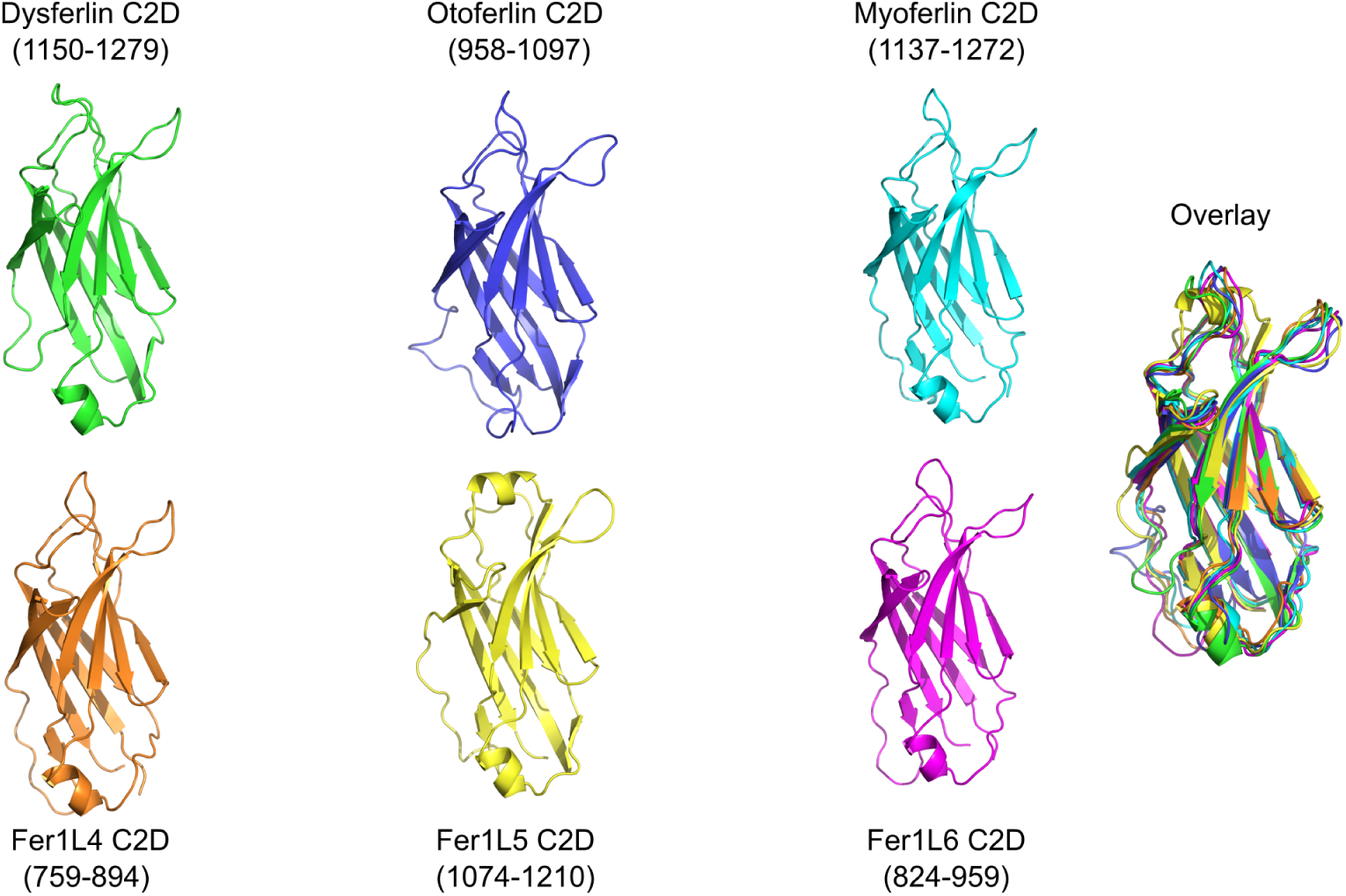
C2D models for dysferlin C2D (green), otoferlin C2D (blue), myoferlin C2D (cyan), Fer1L3 C2D (orange), Fer1L5 C2D (yellow), Fer1L6 C2D (magenta). The superposition of all six C2D models are labeled as ‘Overlay’.

Loop 1 of Fer1L5 has been modeled as a C2C-like *α*-helix. However, unlike the C2C domains in other ferlin proteins, the loop 1 *α*-helix in Fer1L5 is not strongly amphipathic. Fer1L5 Loop 1 does possess two hydrophobic residues that could be used to bind phospholipid membranes. In contrast to the *β*1-2 subdomain in C2C, the Fer1L5 C2D loop 1 helix does not possess residues that could participate in Ca^2+^ coordination. Loop 3 of Fer1L5 also possesses two bulky hydrophobic residues with which to interact with the phospholipid membrane.

### Analysis of ferlin C2E domains

The Type-2 ferlins, otoferlin, Fer1L4, and Fer1L6, were originally thought to lack a C2E domain [51]. As a consequence of the missing C2 domain, an evolutionary analysis of the ferlin proteins as a family required a large gap to align the seven C2 domains of the Type-1 ferlins with the five or six C2 domains of the Type-2 ferlins [51]. The missing C2 domain in the Type-2 ferlins caused a discrepancy in the sequential nomenclature of the ferlin C2 domains. To maintain a uniform nomenclature, all of the ferlin C2 domains were consecutively labeled C2A-C2F, except in Type-1 ferlins where the domain after C2D was labeled as “C2DE”, to reflect its location in the protein relative to its flanking C2 domains [51]. However, it is now clear from the RoseTTAFold and AlphaFold2 models that C2E is indeed present in all ferlins, including the Type-2 ferlins (**Fig. 1**). In light of the discovery of the C2E domain in Type-2 ferlins, we recommend a return to the consecutive C2A-C2G nomenclature for all ferlins, with the exception of Fer1L4 and Fer1L6 that lack a C2A domain (**Fig. 1**).

The ferlin C2E domain contains a large subdomain between *β*-strand 6 and 7 (*β*6-7) (**Fig. 9**). The C2E *β*6-7 subdomain across the ferlin family is generally negatively charged, and it has been modeled as mostly *α*-helical. The overall length of C2E *β*6-7 varies widely between the ferlin proteins, and it is probably the reason C2E was overlooked in the original ferlin family analysis. The longest C2E *β*6-7 subdomains occur within otoferlin and Fer1L6, which are 203 and 214 residues in length, making the subdomains longer than the prototypical C2 domain. The *β*6-7 subdomains in dysferlin, myoferlin, Fer1L4, and Fer1L5 are: 103, 106, 163, and 101 residues in length, respectively. Interestingly, the calpain cleavage site that produces a “synaptotagmin-like” mini-dysferlin_C72_ (C2F-C2G-TM fragment) occurs within the *β*6-7 subdomain of C2E, but only in the exon40a insertion of human dysferlin [70]. Proteolytic cleavage at that site with calpain would remove six *β* strands from the C2E domain and leave two *β* strands as remnants on the N-terminus of the mini-dysferlin_C72_ fragment.

**Fig 9.**
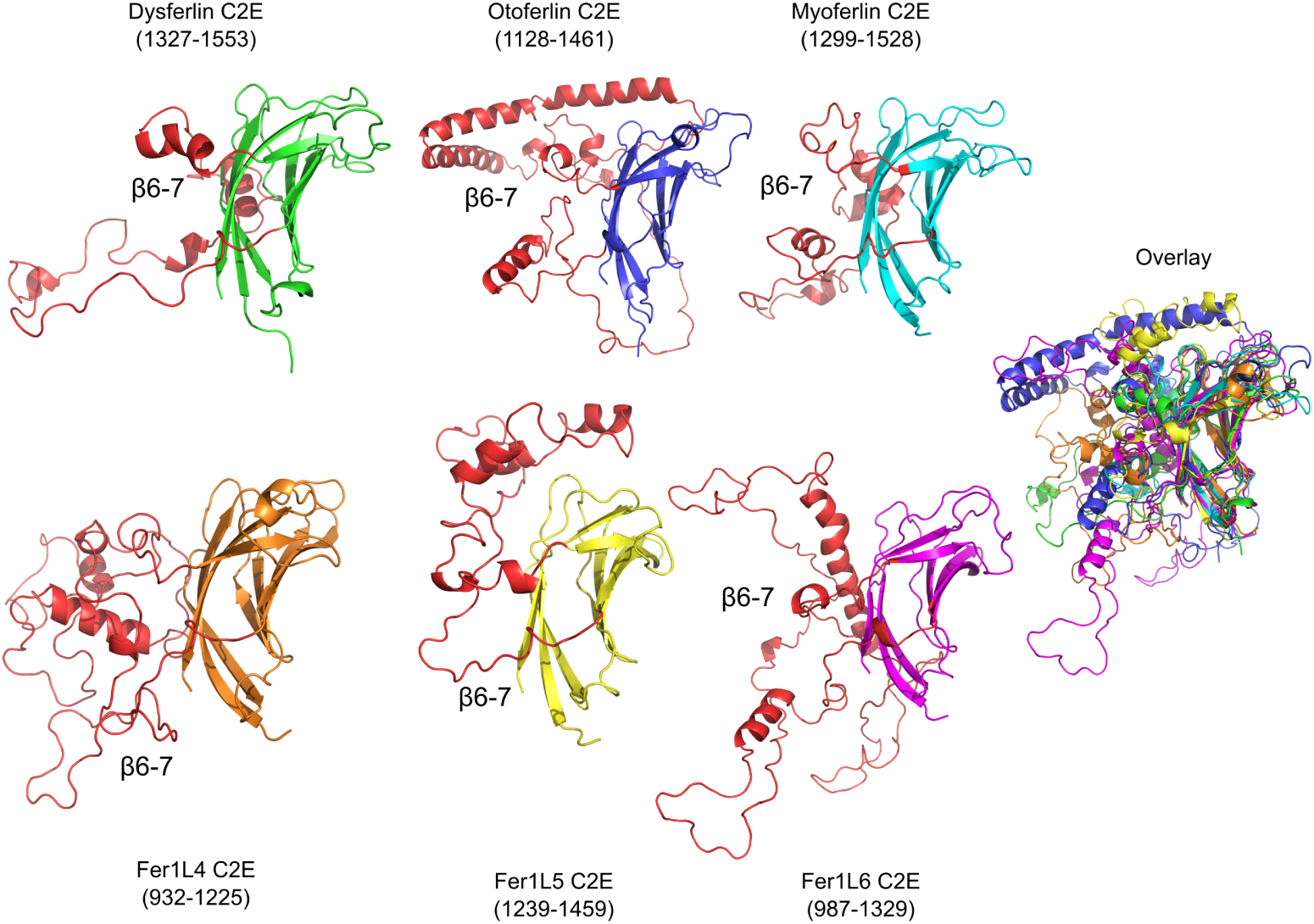
C2E models for dysferlin C2E (green), otoferlin C2E (blue), myoferlin C2E (cyan), Fer1L3 C2E (orange), Fer1L5 C2E (yellow), Fer1L6 C2E (magenta). The *β*6-7 subdomain of C2E is labeled and colored in red. The superposition of all six C2E models are labeled as ‘Overlay’.

It is possible that C2E binds Ca^2+^ in the divalent binding pocket; however, the requisite constellation of acidic residues normally used to coordinate Ca^2+^ is currently unclear. In each of the ferlin C2E domains, there are conserved Arg residues within the divalent cation binding pocket. For example, dysferlin Arg-1409 (**Fig. S7**) may act in the same manner as was observed in the C2Av1 (C2A variant 1) where positively charged residues salt-bridge to the Ca^2+^ binding residues; thus preventing Ca^2+^ binding [52]. The only exception is Fer1L5 where a Trp residue is predicted at that locus.

### Analysis of ferlin C2F domains

The C2F domain also resembles a prototypical C2 domain (**Fig. 10**), with the exception of a unique subdomain between *β* strands 6 and 7(**Fig. S8**). Our ferlin models indicate that C2F possesses all conserved acidic amino acids to bind Ca^2+^, and it likely interacts with phospholipid membranes similar to dysferlin C2A [71]. The ~ 60 amino acid C2F *β*6-7 subdomain adopts an *α*-*β*-*β*-*β*-*α* fold in which the two *α*-helices lie on one face of a three-stranded anti-parallel *β*-sheet. Interestingly, the C2F *β*6-7 subdomain shares structural similarity to a family of double-stranded RNA-binding (dsRNA) domains (**Fig. 11**). For example, the dsRBD5 of the *Drosophila* Staufen protein [72] (1STU) superimposes with the human dysferlin *β*6-7 subdomain of C2F with an RMSD of 2.7Å (**Fig. 11A**), and the RISC-loading complex subunit TARBP2 [73] (4WYQ) superimposes with an RMSD value of 0.86Å (**Fig. 11B**). It is unknown if any of the ferlins can bind dsRNA through the *β*6-7 subdomains of C2F, but the similarity to known dsRNA binding domains is intriguing. The residues that are used for dsRNA binding are not conserved in the ferlin C2F domains; however, otoferlin C2F conserves at least two of the four known residues that bind dsRNA.

**Fig 10.**
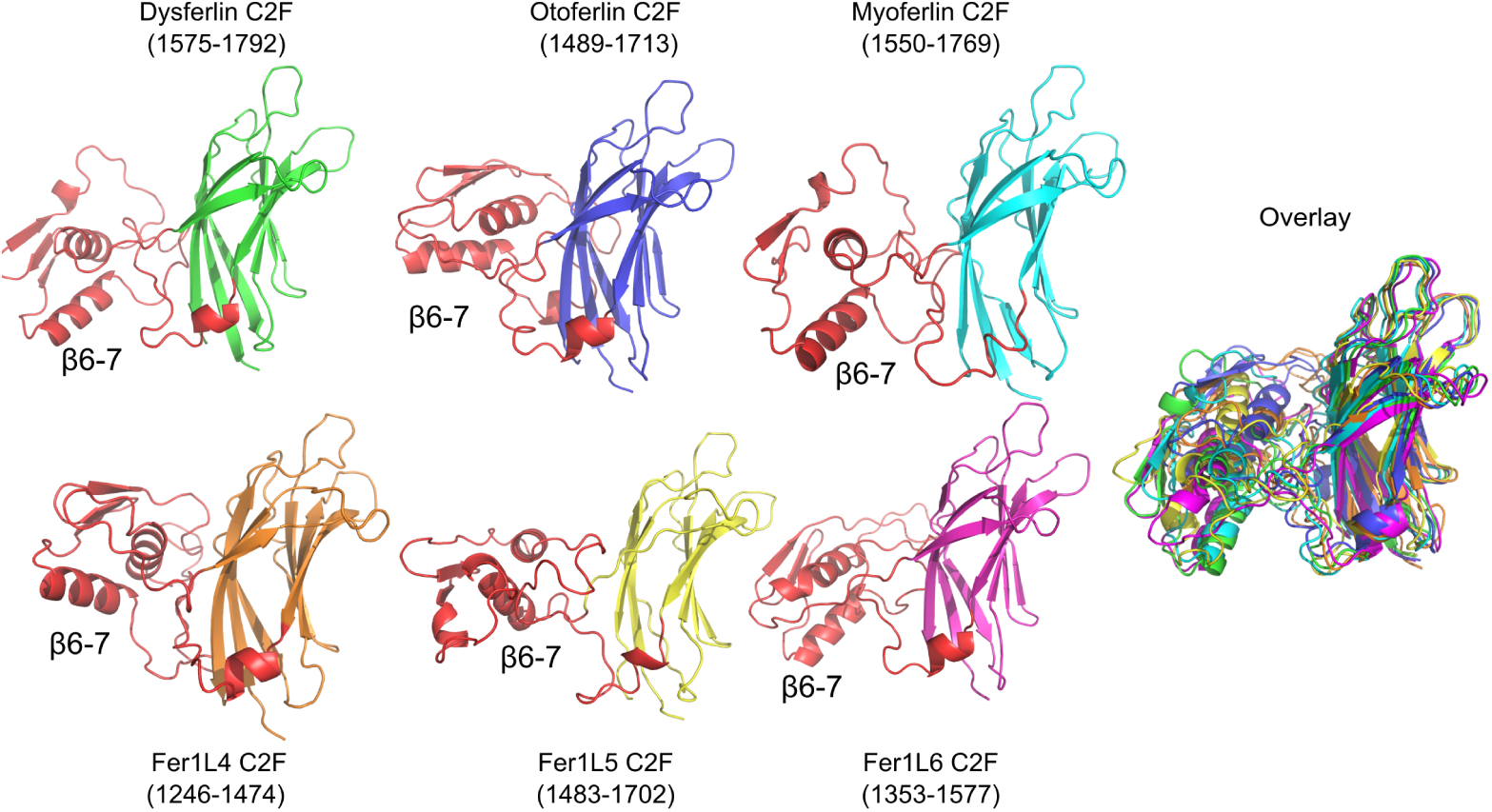
Dysferlin C2F (green), otoferlin C2F (blue), myoferlin C2F (cyan), Fer1L4 C2F (orange), Fer1L5 C2F (yellow), Fer1L6 (magenta). The *β*6-7 subdomain of C2F is labeled and colored red. The overlay of each domain is to the right of the figure.

**Fig 11.**
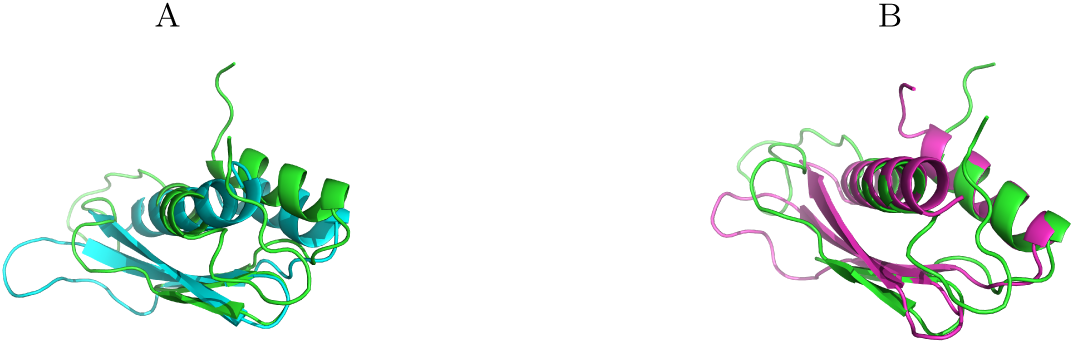
Dysferlin C2F *β*6-7 subdomain aligned with known dsRNA binding domains. **A.** Superimposed *β*6-7 subdomain of Dysferlin C2F (green) and RNA binding domain of Staufen (1STU) in cyan; RMSD = 2.7 Å . **B.** Superimposed *β*6-7 subdomain of Dysferlin C2F (green) and dsRNA binding domain of TARBP2 (4WYQ) in magenta; RMSD = 0.86 Å .

The Type-2 ferlin (otoferlin, Fer1L4, and Fer1L6) C2F domains possess a conserved Lys residue at a critical site on Loop 2 that is typically used for Ca^2+^ coordination in other C2 domains and would likely negatively impact Ca^2+^ binding. The homologous locus in the Type-1 ferlins have residues more consistent with canonical Ca^2+^ coordination in C2 domains. The sequences for myoferlin C2F and Fer1L5 C2F possess the typically Asn residue at this locus; therefore, they are likely to bind Ca^2+^ in a manner similar to C2A. Interestingly, dysferlin C2F places a cysteine residue at this locus; however, the Metal Ion-Binding Site Prediction and Docking Server (MIB) predicts Ca^2+^ binding at the consensus divalent cation binding pocket of C2F [69].

### Analysis of ferlin C2G domains

The C2G domain is the final C2 domain in the ferlin proteins prior to the transmembrane span (**Fig. 1**). The residues that coordinate Ca^2+^ are present within each ferlin C2G domain; therefore, we predict that C2G should bind calcium ions. The models of each C2G domain from all human ferlins are predicted to be remarkably similar despite differences in sequence (**Fig. 12**). Each C2G model superimposes with a mean RMSD of 1.2 Å. There are two predicted subdomains in C2G. The first C2G subdomain has an antiparallel two-stranded *β*-sheet inserted between *β* strands 4 and 5 (*β*4-5). The *β*4-5 subdomain of C2G gives the domain an overall ‘L’ shape. Due to the conservation of large hydrophobic residues at the tip of the C2G *β*4-5 subdomain throughout all ferlins (**Fig. S9**), it is possible that the C2G *β*4-5 subdomain assists in membrane binding. In this model, the *β*4-5 subdomain and loops 1 and 3 of the C2G domain can simultaneously interact with phospholipid membranes to position the C2G domain parallel to the membrane in a 3-point stance. The second subdomain of C2G is located between *β*6-7 and is roughly 36 residues in length across all ferlins. The *β*6-7 subdomain is also well conserved among all ferlins except for Fer1L4 (**Fig. S9**. In Fer1L4, the *β*6-7 subdomain includes two long oppositely charged *α*-helices. The function of the *β*6-7 subdomain is unknown.

**Fig 12.**
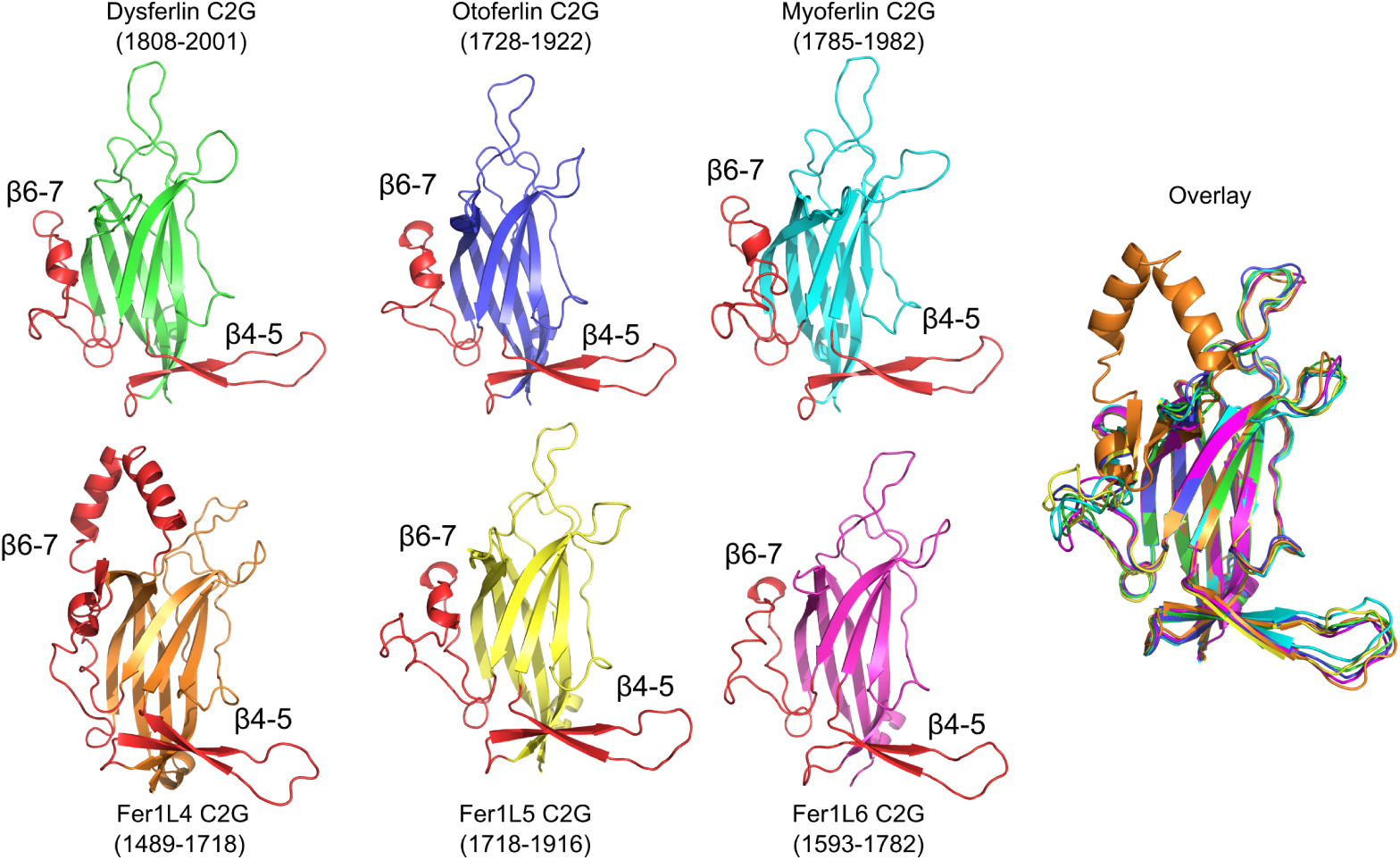
C2G models for dysferlin C2G (green), otoferlin C2G (blue), myoferlin C2G (cyan), Fer1L3 C2G (orange), Fer1L5 C2G (yellow), Fer1L6 C2G (magenta). The *β*4-5 and *β*6-7 subdomains of C2G are labeled and colored red. The superposition of all six C2G models are labeled as ‘Overlay’.

### Transmembrane (TM) spans and Cytoplasmic-Facing (CF) residues

Transmembrane (TM) spans were predicted from the full-length RoseTTAFold ferlin models and confirmed with the TMHMM server [74] (**1**). Each transmembrane span of the ferlins is capped with either a Trp or an Arg on the N-terminal end of the transmembrane span. The transmembrane span for most of the ferlins is predicted to be 22 residues in length except for Fer1L5 that has a 19 amino acid TM (**Fig. S10**). All ferlin transmembrane helices appear to possess two short *α*-helical segments on either side of the TM helix that are gapped by conserved residues. The gapping residues tend to break the helical propensity of the TM. On the N-terminal side, there is a ‘WRRF’ sequence motif in dysferlin where the W and the F residues form the first gap. These residues have previously been implicated in aggregating phosphotidylserine near the TM span [75]. The C-terminal short *α*-helix, potentially extracellular, is gapped by conserved Pro residue that bends or kinks the TM (**Fig. S10**, 1). Indeed, there is a clinically isolated mutation of the conserved Pro residue in otoferlin (P1987R) that is linked to auditory neuropathy [76]. Otoferlin is unusual in that its predicted TM seems to be shifted by the presence of Lys-1968. The shift in the transmembrane prediction places the conserved proline within the membrane and leaves five cytoplasmic-facing residues on the outside of the cell. In dysferlin, there is a clinically-defined transmembrane mutation A2066T that has been linked to LGMD-2B [77]. Ala-2066 occurs in the unstructured gapped residues between the TM and the cytoplasmic-facing helix and could interfere with the overall architecture of the TM segment.

## Discussion

Ferlins were first identified in 1980 as a gene (*fer-1* ) essential for fertilization in *C. elegans* [78]. Later the fer-1 protein was found to regulate Ca^2+^-dependent membrane fusion of membranous organelles during spermatogenesis [79]). A mammalian fer-1 ortholog was later designated as “*dys* -ferlin” to reflect the deleterious role that mutations within the *DYSF* gene play in Limb-Girdle Muscular Dystrophy-2B (LGMD-2B) [80]. Five additional *fer-1* paralogs were subsequently annotated as ferlins in mammalian genomes. Phylogenetic analysis revealed that the ferlin family likely has an ancestral function as regulators of vesicle fusion and receptor trafficking [51, 81]. In the last several years, the role of ferlins has expanded to include vesicle fusion and plasma membrane repair [2, 9, 19, 82]. However, despite the indirect evidence implicating ferlins in membrane fusion, the mechanism is currently unknown.

The multi-domain nature and the inherent flexibility of ferlins have limited the biophysical analysis of the full-length ferlin proteins. One approach towards understanding ferlin structure has been to methodically deconstruct the proteins into their respective protein domains and study the 3D structure of each isolated domain, in addition to each domain’s functional contribution to the activity of the whole protein. This piecemeal strategy has been successful in determining the structures for the C2A domains of dysferlin [52, 54], myoferlin [53, 56], and otoferlin [55], as each has been solved by X-ray crystallography or NMR spectroscopy. Further, the inner DysF domain structure of dysferlin and myoferlin have also been solved using both techniques [62, 63]. The boundaries of the ferlin domains that have been successfully analyzed to date have been fairly straightforward to determine, but the remaining domains have proved to be more difficult. Therefore, determining accurate boundaries for the remaining structural aspects of the ferlins will be useful to isolate stable domains that are amenable for analysis with biophysics techniques. The amino acid-level precision required to predict stable domains is provided by the RoseTTAFold and Alphafold2 algorithms.

It is interesting to note the C2 domain loops that have been extended in the ferlins as conserved subdomains. C2A, C2B, and C2D do not possess extraneous subdomains, while C2C, C2-FerA, C2E, C2F, and C2G have elongated loops with unique characteristics. The biological functions of some of the ferlin C2 subdomains can be postulated based on their similarity to structurally homologous proteins. For example, C2C is the only C2 domain in the ferlins that includes subdomains within the putative Ca^2+^/phospholipid interface of the domain (**Fig. 13**). The *β*1-2 loop in C2C likely adds an additional hydrophobic surface to its Ca^2+^-binding pocket with which to interact with phospholipids. A similar, but smaller, loop insertion is present within phospholipase A2 [83]. The functions of the additional two C2C subdomains are not clear. Next, the subdomain between *β*4-5 in C2-FerA is composed of a four-helix bundle (FerA), which projects out from the domain (**Fig. 5**). The FerA subdomain in isolation has been implicated in Ca^2+^-dependent membrane binding although the mechanism has not been described [61]. From the model of C2-FerA, the domain likely binds phospholipids through hydrophobic interactions between with the “opened” FerA subdomain and a compatible membrane surface, but the body of its C2 domain does not possess the Ca^2+^-binding residues or the phospholipid binding residues for typical C2 domain membrane binding. The C2E domain is the next highly embellished C2 domain in the ferlins. The *β*6-7 loop of C2E is at least as large as the domain itself, but the physiological function of such a large insertion is difficult to assess. It is possible that the sheer size of the C2E *β*6-7 subdomain and the overall negative charge of the subdomain provides a mechanism to join other domains throughout the ferlin protein. Such an extensive interaction may serve to restrain the overall shape of the full-length molecule. The loop insertion for C2F across all ferlins is particularly interesting. C2F’s *β*6-7 insertion shares similarity with double-stranded RNA binding motifs. If this domain does indeed bind dsRNA, it is not yet clear how any future ferlin dsRNA-binding activity relates to membrane interaction. Lastly, the *β*4-5 subdomain of C2G gives its C2 domain an overall ‘L’ shape. The large hydrophobic residues of *β*4-5 likely contribute to membrane binding. Indeed, this property of *β*4-5 of C2G has been conserved across all six ferlins (**Fig. S9**). In total, most of the C2 domain loop structures in the ferlins have been enhanced, with the exception of loops 2 and 3 that typically form part of the Ca^2+^ binding pocket (**Fig. 13**). The placement of subdomains in the various ferlin C2 domains suggests that Ca^2+^-dependent phospholipid-binding may be a protected activity.

**Fig 13.**
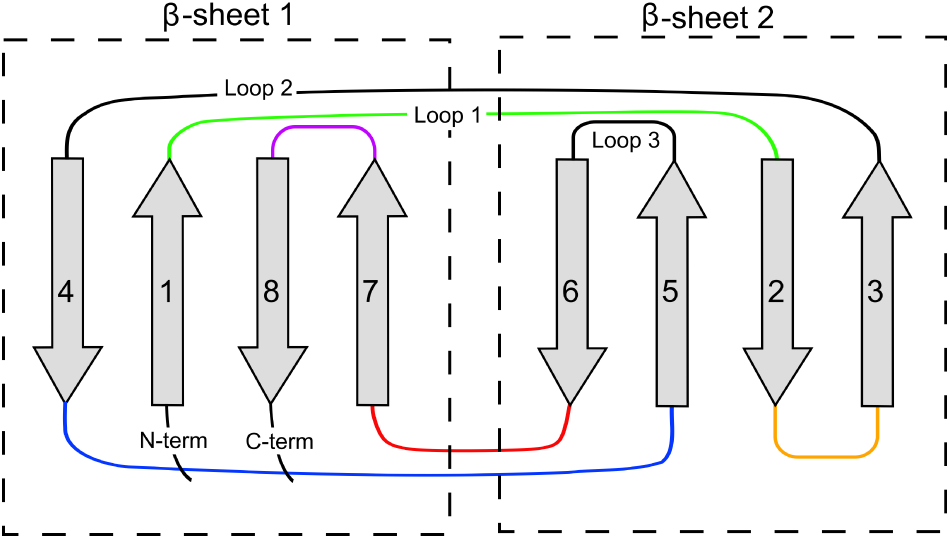
Schematic of Type-II C2 domain. The schematic highlights the loops with embedded subdomains in the various ferlin C2 domains. The Type-2 C2 domain *β*-strand topology is shown as grey arrows. The colored loops: *β*1-2 (green, C2C), *β*2-3 (orange, C2-FerA), *β*4-5 (blue, C2-FerA, C2G), *β*6-7 (red, C2C, C2E, C2G, C2F), and *β*7-8 (purple, C2C) show where conserved subdomains are present.

Our RoseTTAFold and AlphaFold2 results provide an excellent tool to decipher the domain structure of the ferlins, but further questions remain. For example, what are the ligand binding residues of each domain, how do ferlins associate with membranes, how do pathogenic mutations affect the function of the protein, what is the function of the various subdomains, and does domain-domain intramolecular interactions contribute to the overall activity? Understanding the structures and molecular mechanisms of large multifunctional ferlin proteins will be an important step towards understanding and treating ferlin-associated diseases such as Limb-Girdle Muscular Dystrophy type-2B (dysferlin), auditory neuropathy (otoferlin) and, and perhaps in the treatment of metastatic tumors (myoferlin).

## Acknowledgments

The authors would like to thank the Jain Foundation for their support of this work. The authors would also like to acknowledge Drs. Nick Grishin and Jimin Pei for the initial evidence of our C2-FerA model.

## Author contributions

Matthew J. Dominguez, conceptualization, data curation, formal analysis, writing-original draft; Jon J. McCord, data curation; R. Bryan Sutton, conceptualization, data curation, formal analysis, writing-original draft, supervision, funding aquisition.

## Data availability

All of the extracted domains from all six human ferlins are available as pdb files at the following address: 10.6084/m9.figshare.17912222

## Supporting information

**Table 1.** Predicted Ferlin Transmembrane domain boundaries and Cytoplasmic-facing residues. The conserved ferlin Pro residues are listed.

|  | Transmembrane Span | Conserved Proline | Cytoplasmic-Facing |
| --- | --- | --- | --- |
| Dysferlin (Fer1L1) | 2045 - 2067 | Pro-2068 | 2068 - 2080 (12) |
| Otoferlin (Fer1L2) | 1969 - 1991 | Pro-1987 | 1992 - 1997 (5) |
| Myoferlin (Fer1L3) | 2026 - 2046 | Pro-2049 | 2048 - 2061 (13) |
| Fer1L4 | 1758 - 1780 | Pro-1781 | 1781 - 1794 (13) |
| Fer1L5 | 1962 - 1981 | Pro-1983 | 1982 - 2057 (75) |
| Fer1L6 | 1824 - 1846 | Pro-1846 | 1847 - 1857 (10) |

**Table S1.** Model Scores for Dysferlin domains

| Dysferlin | C2A | C2B | C2C | C2-FerA | DysF | C2D | C2E | C2F | C2G |
| --- | --- | --- | --- | --- | --- | --- | --- | --- | --- |
| MolProbity Score | 1.46 | 1.40 | 1.44 | 1.48 | 1.92 | 1.84 | 1.18 | 1.30 | 1.37 |
| Clash Score | 0.52 | 0.99 | 0 | 0.46 | 1.61 | 0.99 | 0 | 0.58 | 0.32 |
| Ramachandran Favored (%) | 89.52 | 93.40 | 87.12 | 90.04 | 92.08 | 92.31 | 90.62 | 96.02 | 93.08 |
| Ramachandran Outliers (%) | 0.95 | 0 | 3.07 | 3.03 | 2.08 | 0.96 | 2.60 | 0.57 | 0.63 |
| QMEAN (Z-score) | -0.64 | -0.87 | -2.75 | -2.21 | -3.76 | -1.65 | -3.71 | -1.13 | -2.78 |

**Table S2.** Model Scores for Otoferlin domains

| Otoferlin | C2A | C2B | C2C | C2-FerA | C2D | C2E | C2F | C2G |
| --- | --- | --- | --- | --- | --- | --- | --- | --- |
| MolProbity Score | 1.44 | 1.34 | 1.18 | 1.41 | 1.29 | 1.42 | 1.59 | 1.51 |
| Clash Score | 2.11 | 1.53 | 0.00 | 0.58 | 0.46 | 0.00 | 0.87 | 1.62 |
| Ramachandran Favored (%) | 92.86 | 93.07 | 90.21 | 93.17 | 93.39 | 90.39 | 90.12 | 90.91 |
| Ramachandran Outliers (%) | 0.00 | 1.98 | 3.50 | 0.72 | 3.31 | 3.16 | 0.58 | 1.82 |
| QMEAN (Z-score) | -1.09 | -1.07 | -3.43 | -1.66 | -2.57 | -3.34 | -1.26 | -2.54 |

**Table S3.**
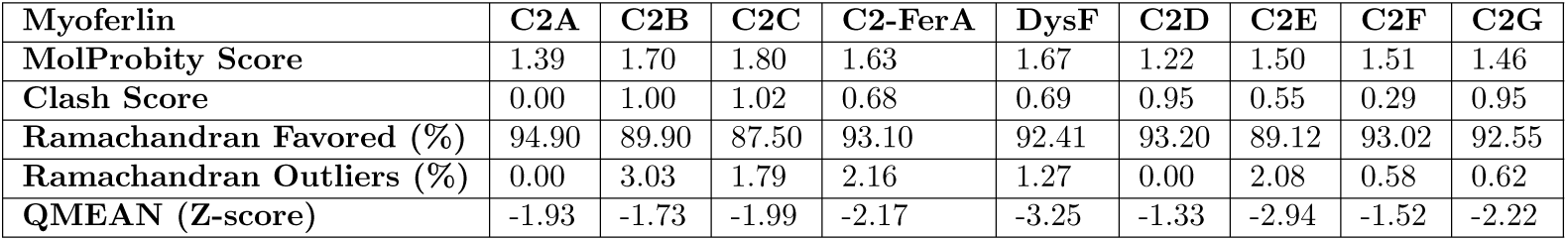
Model Scores for Myoferlin domains

**Table S4.** Model Scores for Fer1L4 domains

| **Fer1L4** | **C2B** | **C2C** | **C2-FerA** | **C2D** | **C2E** | **C2F** | **C2G** |
| --- | --- | --- | --- | --- | --- | --- | --- |
| **MolProbity Score** | 1.70 | 1.60 | 1.52 | 1.57 | 1.52 | 1.96 | 1.33 |
| **Clash Score** | 1.46 | 1.57 | 0.62 | 0.93 | 0.22 | 0.58 | 0.54 |
| **Ramachandran Favored (%)** | 92.59 | 89.93 | 91.04 | 88.6 | 90.73 | 83.33 | 91.51 |
| **Ramachandran Outliers (%)** | 0.93 | 2.68 | 1.43 | 0.88 | 2.02 | 4.30 | 1.89 |
| **QMEAN (Z-score)** | -1.71 | -2.69 | -1.46 | -1.05 | -3.56 | -2.94 | -1.97 |

**Table S5.** Model Scores for Fer1L5 domains

| **Fer1L5** | **C2A** | **C2B** | **C2C** | **C2-FerA** | **DysF** | **C2D** | **C2E** | **C2F** | **C2G** |
| --- | --- | --- | --- | --- | --- | --- | --- | --- | --- |
| **MolProbity Score** | 1.74 | 1.52 | 1.66 | 1.37 | 1.72 | 1.57 | 1.91 | 1.42 | 2.06 |
| **Clash Score** | 0.50 | 0.50 | 0.41 | 0.23 | 0.65 | 1.35 | 1.40 | 0.29 | 1.56 |
| **Ramachandran Favored (%)** | 89.32 | 92.31 | 81.20 | 92.41 | 89.11 | 90.91 | 87.43 | 89.67 | 89.51 |
| **Ramachandran Outliers (%)** | 1.94 | 0.00 | 2.56 | 0.84 | 2.42 | 2.73 | 1.57 | 1.09 | 1.23 |
| **QMEAN (Z-score)** | -1.19 | -1.84 | -1.27 | -2.10 | -5.13 | -2.34 | -2.44 | -1.21 | -1.83 |

**Table S6.**
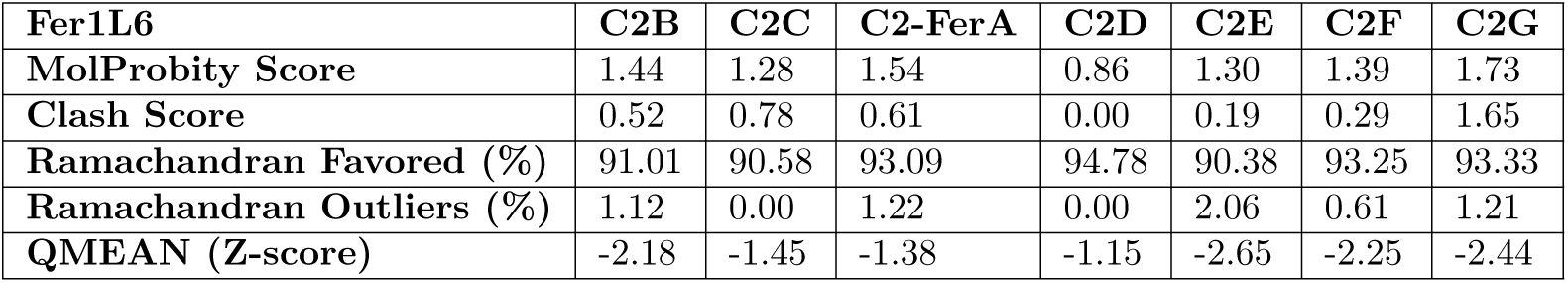
Model Scores for Fer1L6 domains

| **Fer1L6** | **C2B** | **C2C** | **C2-FerA** | **C2D** | **C2E** | **C2F** | **C2G** |
| --- | --- | --- | --- | --- | --- | --- | --- |
| **MolProbity Score** | 1.44 | 1.28 | 1.54 | 0.86 | 1.30 | 1.39 | 1.73 |
| **Clash Score** | 0.52 | 0.78 | 0.61 | 0.00 | 0.19 | 0.29 | 1.65 |
| **Ramachandran Favored (%)** | 91.01 | 90.58 | 93.09 | 94.78 | 90.38 | 93.25 | 93.33 |
| **Ramachandran Outliers (%)** | 1.12 | 0.00 | 1.22 | 0.00 | 2.06 | 0.61 | 1.21 |
| **QMEAN (Z-score)** | -2.18 | -1.45 | -1.38 | -1.15 | -2.65 | -2.25 | -2.44 |

**Fig S1.**
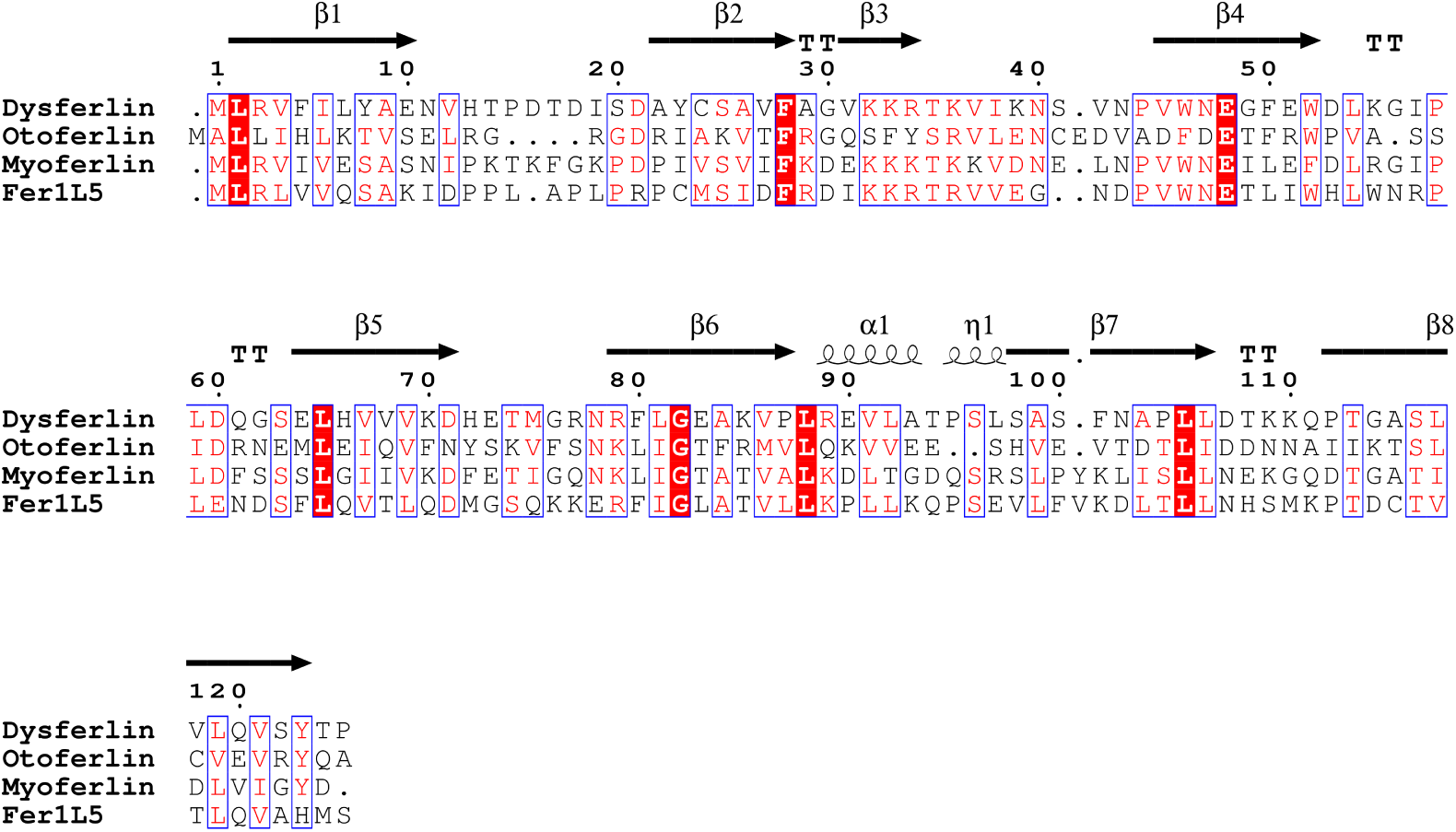
Structure based primary sequence alignment of the C2A domains of the ferlin family. Secondary structure assignments are based on the known and predicted structures of dysferlin as calculated by DSSP. The *η* symbol refers to a 3_10_-helix. *α*-helices, 3_10_-helices and *π*-helices are displayed as medium, small and large squiggles, respectively. *β*-strands are rendered as arrows, strict *β*-turns as TT letters and strict *α*-turns as TTT. Residues that are absolutely conserved between all C2A domains are highlighted in red. Conserved residues are boxed in blue. Numbers along the top of the alignment correspond to residue numbers in the dysferlin sequence. The mean evolutionary relatedness between the C2A domains of dysferlin, otoferlin, myoferlin, and Fer1L5 is 26.5% identity and 56.7% similarity.

**Fig S2.**
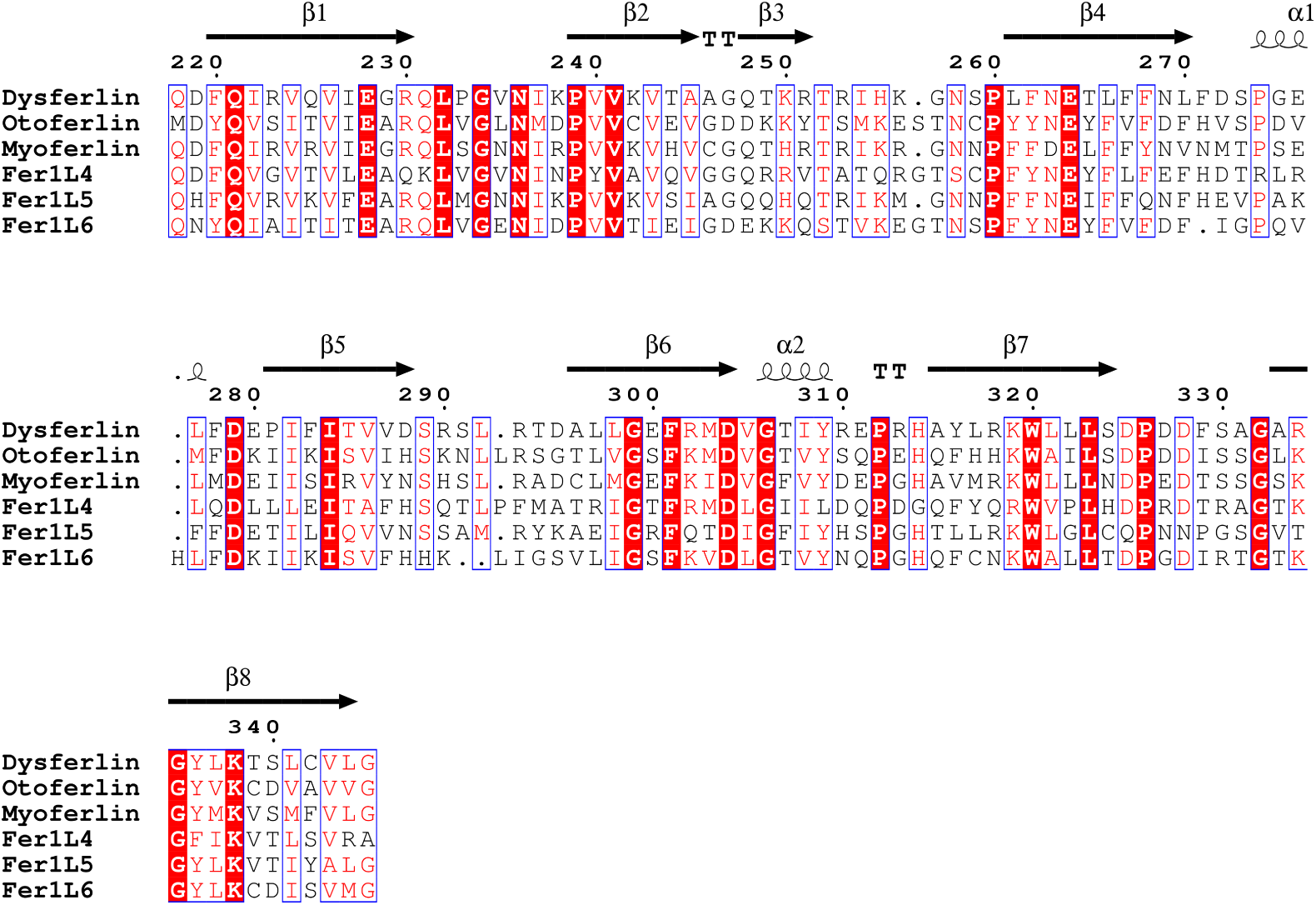
Model-based primary sequence alignment of the C2B domains of the ferlin family. Secondary structure assignments are based on the predicted structures of dysferlin C2B as calculated by DSSP. The *η* symbol refers to a 3_10_-helix. *α*-helices, 3_10_-helices and *π*-helices are displayed as medium, small and large squiggles, respectively. *β*-strands are rendered as arrows, strict *β*-turns as TT letters and strict *α*-turns as TTT. Residues that are absolutely conserved between all C2B domains are highlighted in red. Conserved residues are boxed in blue. Numbers along the top of the alignment correspond to residue numbers in the human dysferlin sequence. The mean evolutionary relatedness between the C2B domains of dysferlin, otoferlin, myoferlin, Fer1L4, Fer1L5, and Fer1L6 is 44.4% mean identity and 63.8% mean sequence similarity across all six C2B domains.

**Fig S3.**
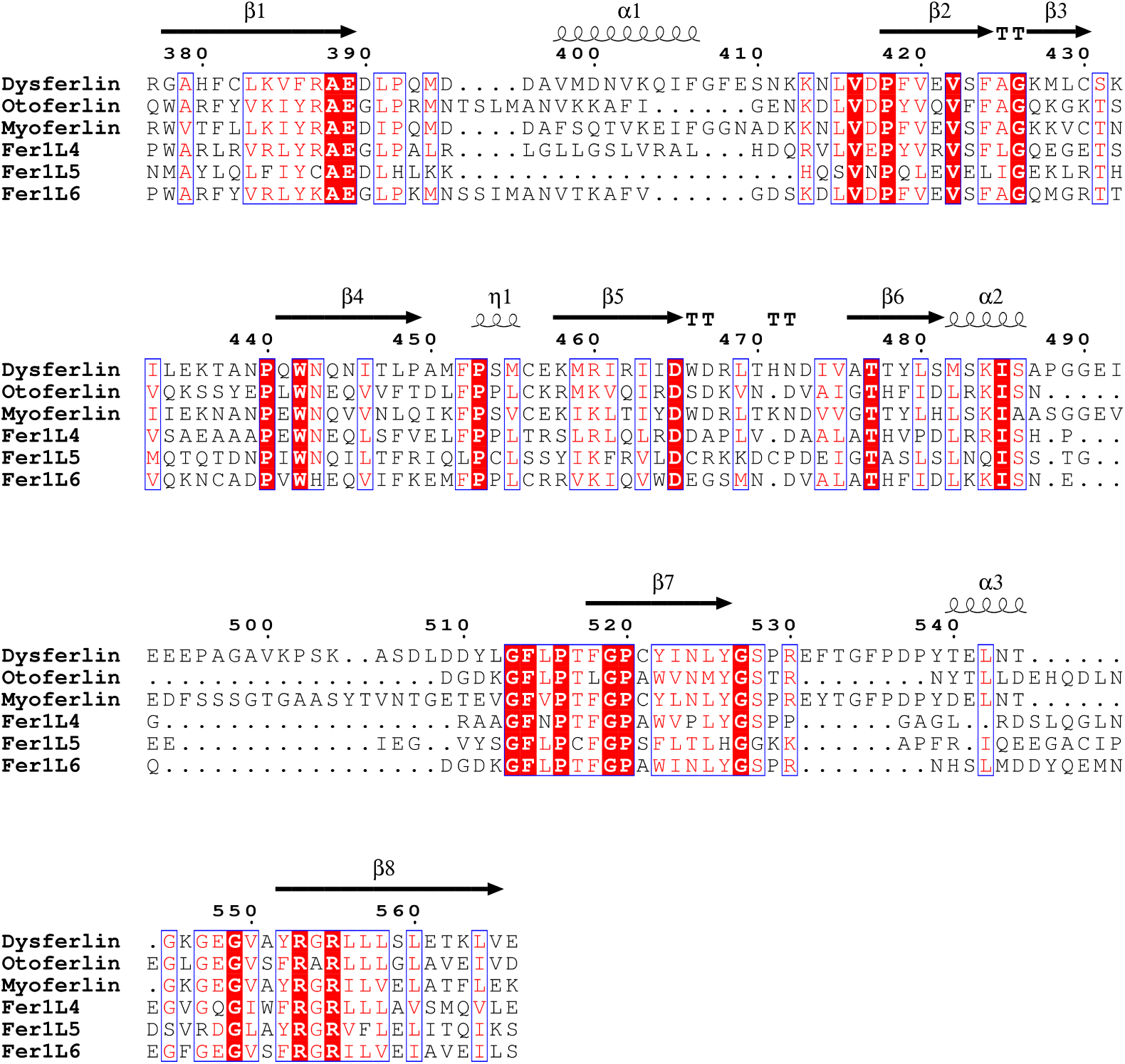
Model-based primary sequence alignment of the C2C domains of the ferlin family. Secondary structure assignments are based on the predicted structures of dysferlin C2C as calculated by DSSP. The *η* symbol refers to a 3_10_-helix. *α*-helices, 3_10_-helices and *π*-helices are displayed as medium, small and large squiggles, respectively. *β*-strands are rendered as arrows, strict *β*-turns as TT letters and strict *α*-turns as TTT. Residues that are absolutely conserved between all C2C domains are highlighted in red. Conserved residues are boxed in blue. Numbers along the top of the alignment correspond to residue numbers in the human dysferlin C2C sequence. The mean evolutionary relatedness between the C2C domains of dysferlin, otoferlin, myoferlin, Fer1L4, Fer1L5, and Fer1L6 is 38.2% mean identity and 60.7% mean sequence similarity.

**Fig S4.**
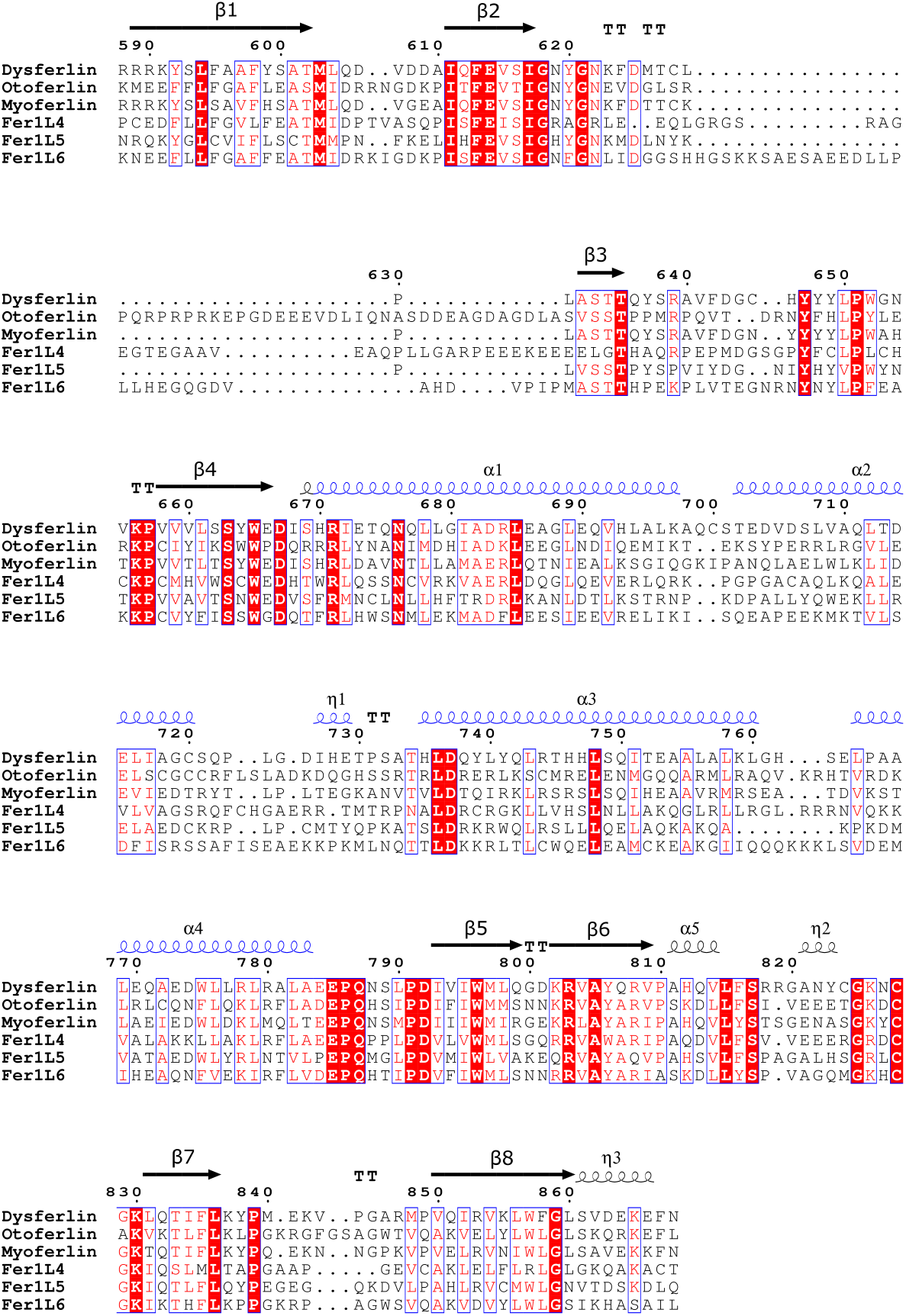
Model-based primary sequence alignment of the C2-FerA domains of the ferlin family. Secondary structure assignments are based on the predicted structures of dysferlin C2-FerA as calculated by DSSP. The *η* symbol refers to a 3_10_-helix. *α*-helices, 3_10_-helices and *π*-helices are displayed as medium, small and large squiggles, respectively. *β*-strands are rendered as arrows, strict *β*-turns as TT letters and strict *α*-turns as TTT. Residues that are absolutely conserved between all C2-FerA domains are highlighted in red. Conserved residues are boxed in blue. Numbers along the top of the alignment correspond to residue numbers in the human dysferlin sequence. The blue helical cartoons demarcate the FerA helices between *β* strand-4 and *β* strand-5. The mean evolutionary relatedness between the C2-FerA domains of dysferlin, otoferlin, myoferlin, Fer1L4, Fer1L5, and Fer1L6 shows 34.9% mean identity and 55.3% mean sequence similarity for all six C2-FerA domains.

**Fig S5.**
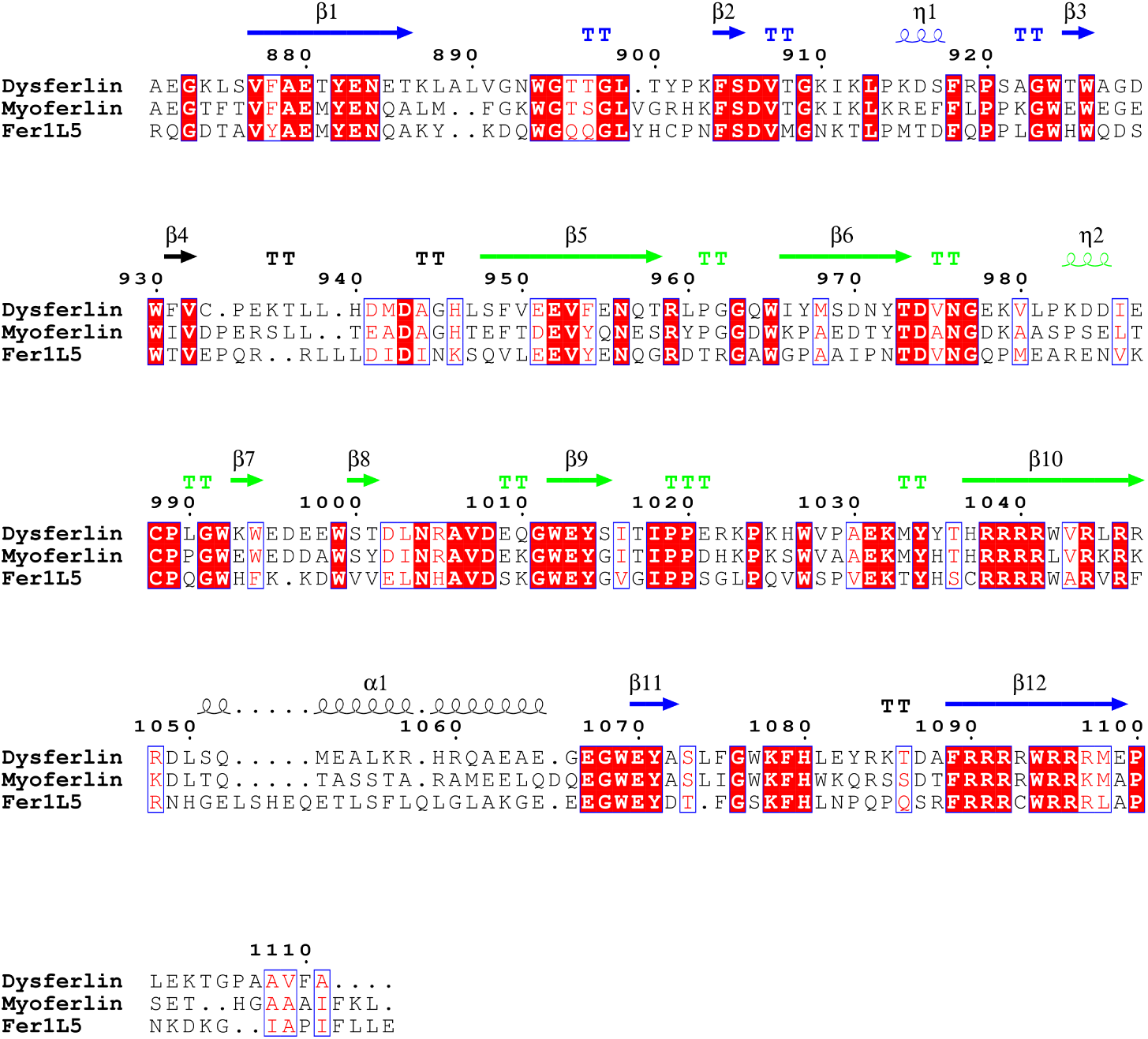
Model-based primary sequence alignment of the DysF domains of the ferlin family. Secondary structure assignments are based on the known and predicted structures of dysferlin DysF as calculated by DSSP. The blue secondary structure is predicted to constitute the “outer” DysF domain, while the green secondary structure demarcates the “inner” DysF domain. The *η* symbol refers to a 3_10_-helix. *α*-helices, 3_10_-helices and *π*-helices are displayed as medium, small and large squiggles, respectively. *β*-strands are rendered as arrows, strict *β*-turns as TT letters and strict *α*-turns as TTT. Residues that are absolutely conserved between all DysF domains are highlighted in red. Conserved residues are boxed in blue. Numbers along the top of the alignment correspond to residue numbers in the human dysferlin sequence. The fold of the predicted dysferlin outer DysF domain superimposes with the known structure of the dysferlin inner DysF structure with an RMSD of 3.5 Å across all C-*α* residues (Figure 7. The mean evolutionary relatedness between the three DysF regions of the Type-I ferlins is 45.0% mean identity and 68.4% mean sequence similarity.

**Fig S6.**
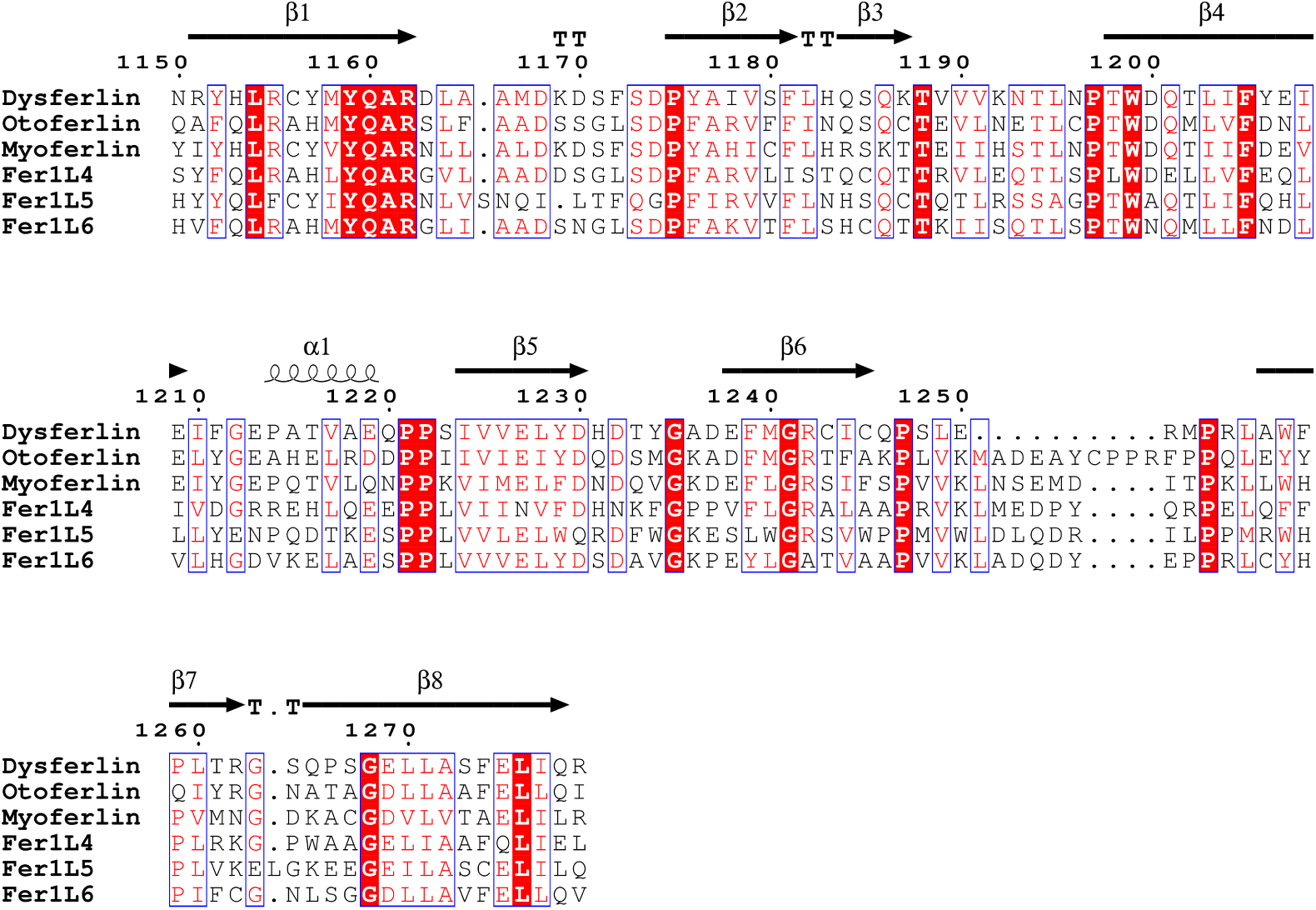
Model-based primary sequence alignment of the C2D domains of the ferlin family. Secondary structure assignments are based on the predicted structures of dysferlin C2D as calculated by DSSP. *α*-helices are displayed as medium squiggles. *β*-strands are rendered as arrows, strict *β*-turns as TT letters. Residues that are absolutely conserved between all C2D domains are highlighted in red. Conserved residues are boxed in blue. Numbers along the top of the alignment correspond to residue numbers in the human dysferlin sequence. The mean evolutionary relatedness between all six C2D domains shows 63.4% mean sequence similarity and 40.9% mean identity.

**Fig S7.**
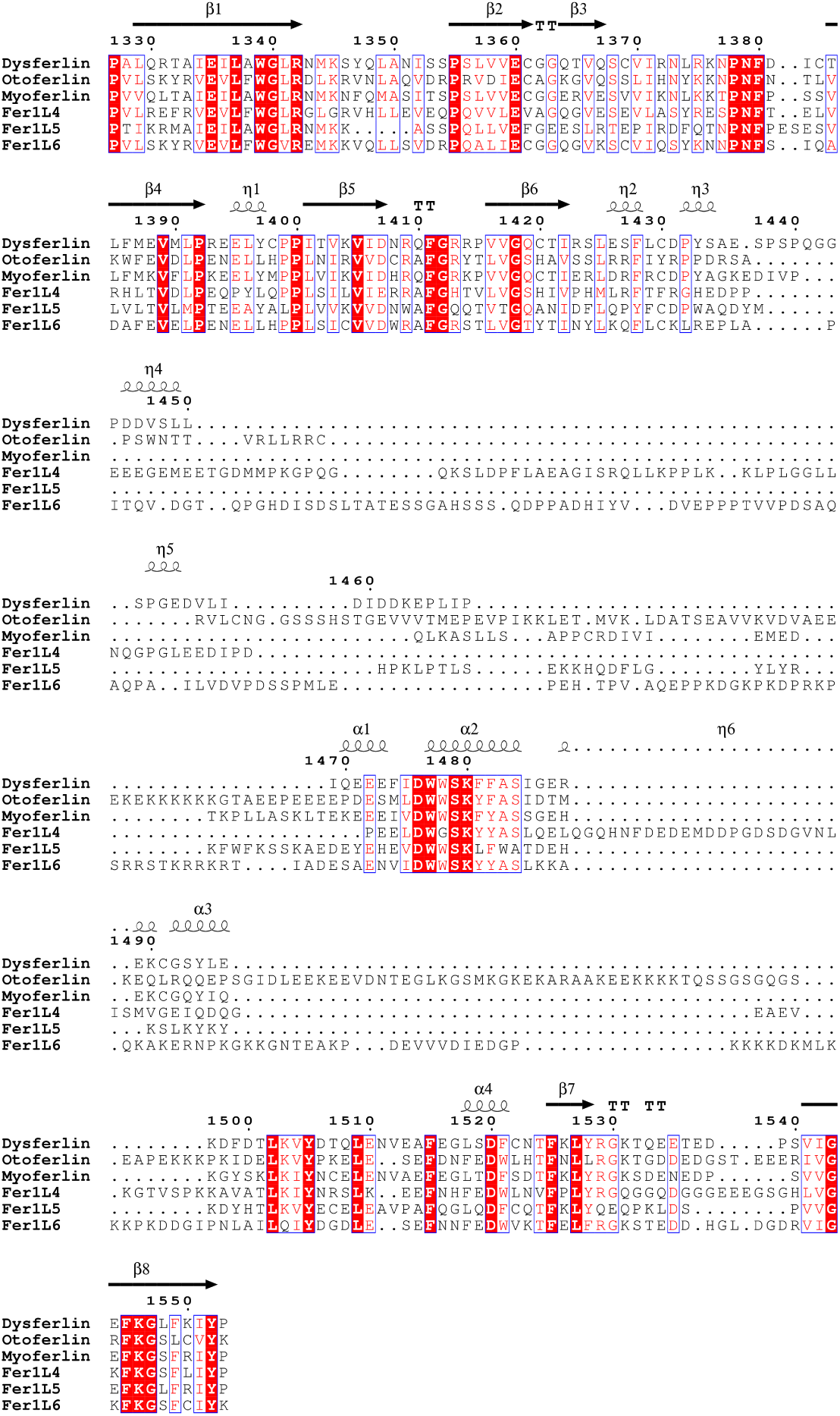
Model-based primary sequence alignment of the C2E domains of the ferlin family. Secondary structure assignments are based on the known and predicted structures of dysferlin C2E as calculated by DSSP. *α*-helices are displayed as medium squiggles. *β*-strands are rendered as arrows, strict *β*-turns as TT letters. Residues that are absolutely conserved between all C2D domains are highlighted in red. Conserved residues are boxed in blue. Numbers along the top of the alignment correspond to residue numbers in the human dysferlin sequence. The mean evolutionary relatedness between the C2E domains of all ferlins is 54.1% mean sequence similarity and 34.7% mean identity.

**Fig S8.**
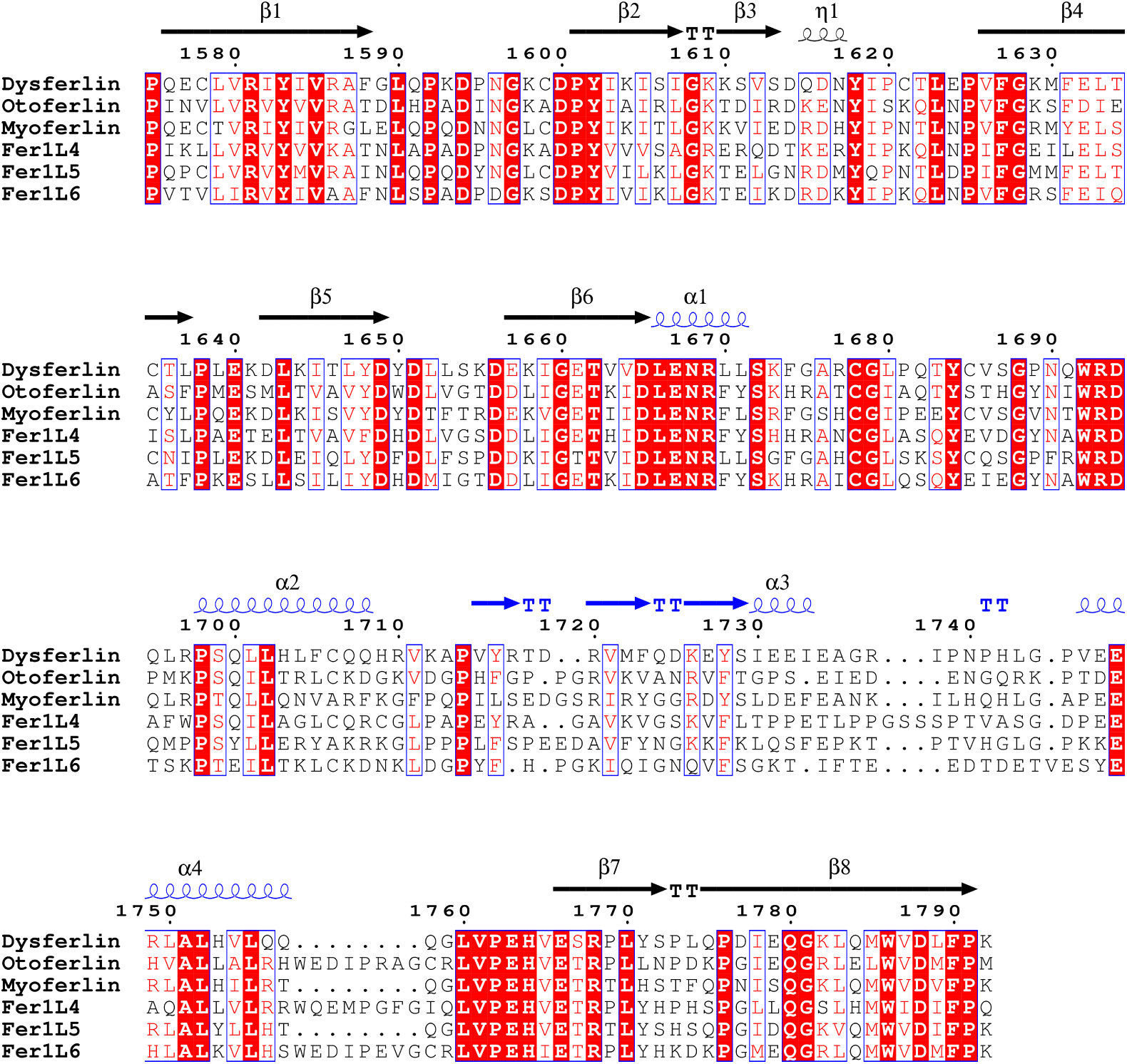
Model-based primary sequence alignment of the C2F domains of the ferlin family. Secondary structure assignments are based on the predicted structures of dysferlin C2F as calculated by DSSP. *α*-helices are displayed as medium squiggles. *β*-strands are rendered as arrows, strict *β*-turns as TT letters. Residues that are absolutely conserved between all C2F domains are highlighted in red. Conserved residues are boxed in blue. Numbers along the top of the alignment correspond to residue numbers in the human dysferlin sequence. beta strands are labeled according to the main body of the C2 domain. The accessory domain between *β*-6 and *β*-7 are colored in blue. The mean evolutionary relatedness between the six C2F domains shows 66.7% mean sequence similarity and 48.2% mean identity for all six C2F domains.

**Fig S9.**
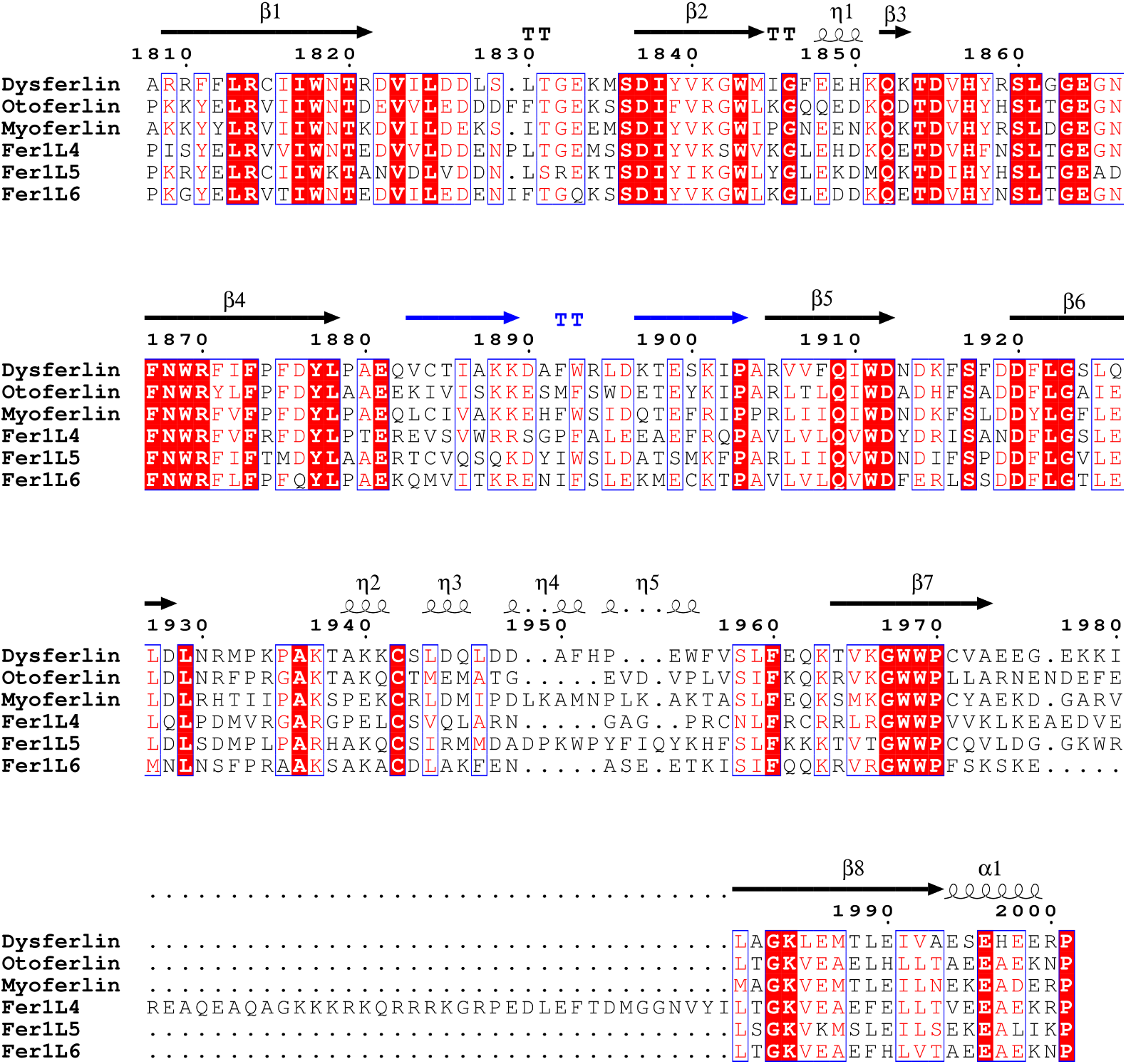
Model-based primary sequence alignment of the C2G domains of the ferlin family. Secondary structure assignments are based on the known and predicted structures of dysferlin C2G as calculated by DSSP. *α*-helices are displayed as medium squiggles. *β*-strands are rendered as arrows, strict *β*-turns as TT letters. Residues that are absolutely conserved between all C2G domains are highlighted in red. Conserved residues are boxed in blue. Numbers along the top of the alignment correspond to residue numbers in the human dysferlin sequence. The mean evolutionary relatedness between the six C2G domains shows 69.5% mean sequence similarity and 51.2% mean identity for all six C2G domains.

**Fig S10.**
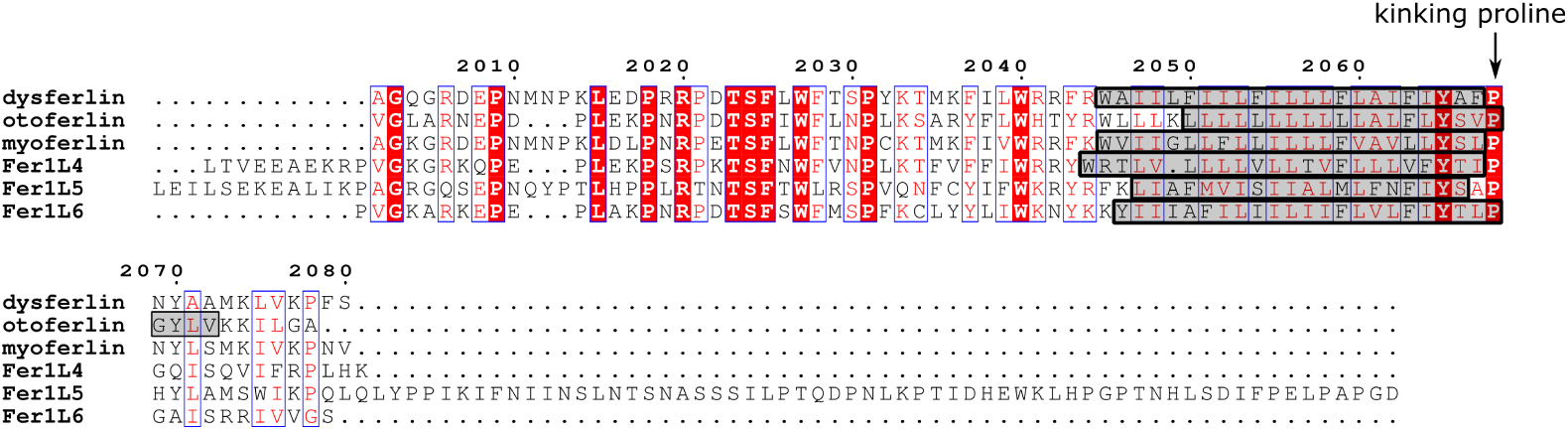
Residues that extend from the C-terminus of C2G to the C-terminus of the ferlin. The predicted transmembrane residues are highlighted in grey.

**Listing 1.**
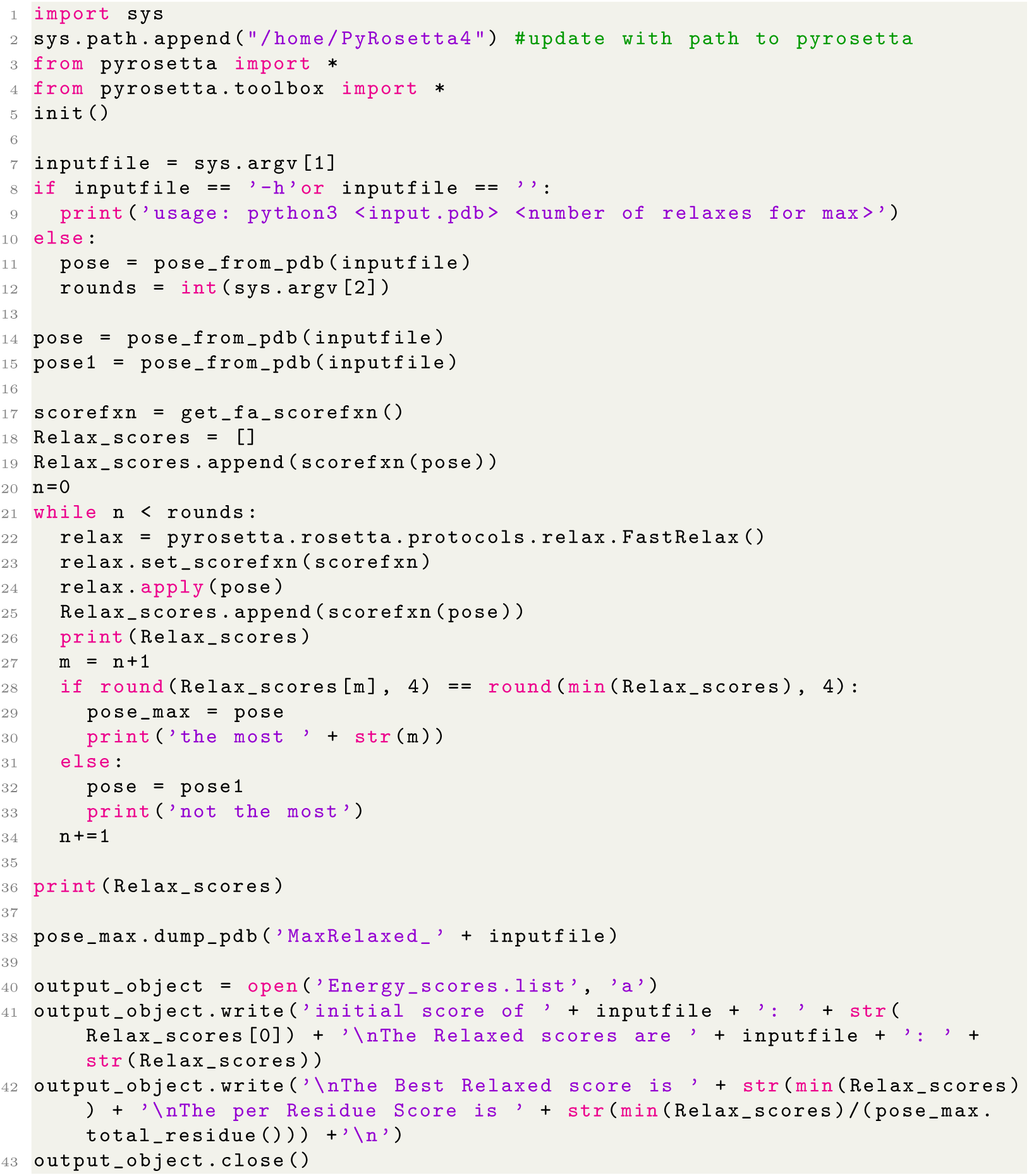
Energy minimization script used to relax

## Notes

https://doi.org/10.6084/m9.figshare.17912222

### Competing Interest Statement

The authors have declared no competing interest.

## References

1. Bulankina AV, Thoms S. Functions of Vertebrate Ferlins. Cells. 2020;9(3):534. doi:10.3390/cells9030534.

2. Bansal D, Campbell KP. Dysferlin and the plasma membrane repair in muscular dystrophy. Trends in Cell Biology. 2004;14(4):206–213. doi:10.1016/j.tcb.2004.03.001.

3. Han R, Campbell KP. Dysferlin and muscle membrane repair. Current Opinion in Cell Biology. 2007;19(4):409–416. doi:10.1016/j.ceb.2007.07.001.

4. Bansal D, Miyake K, Vogel SS, Groh S, Chen CC, Williamson R, et al. Defective membrane repair in dysferlin-deficient muscular dystrophy. Nature. 2003;423(6936):168–172. doi:10.1038/nature01573.

5. Schug N, Braig C, Zimmermann U, Engel J, Winter H, Ruth P, et al. Differential expression of otoferlin in brain, vestibular system, immature and mature cochlea of the rat. European Journal of Neuroscience. 2006;24(12):3372–3380. doi:10.1111/j.1460-9568.2006.05225.x.

6. Takago H, Oshima-Takago T, Moser T. Disruption of Otoferlin Alters the Mode of Exocytosis at the Mouse Inner Hair Cell Ribbon Synapse. Frontiers in Molecular Neuroscience. 2019;11. doi:10.3389/fnmol.2018.00492.

7. Manchanda A, Chatterjee P, Bonventre JA, Haggard DE, Kindt KS, Tanguay RL, et al. Otoferlin Depletion Results in Abnormal Synaptic Ribbons and Altered Intracellular Calcium Levels in Zebrafish. Scientific Reports. 2019;9(1). doi:10.1038/s41598-019-50710-2.

8. Pangršič T, Reisinger E, Moser T. Otoferlin: a multi-C2 domain protein essential for hearing. Trends in Neurosciences. 2012;35(11):671–680. doi:10.1016/j.tins.2012.08.002.

9. Doherty KR, Cave A, Davis DB, Delmonte AJ, Posey A, Earley JU, et al. Normal myoblast fusion requires myoferlin. Development. 2005;132(24):5565–5575. doi:10.1242/dev.02155.

10. Blomme A, Costanza B, de Tullio P, Thiry M, Simaeys GV, Boutry S, et al. Myoferlin regulates cellular lipid metabolism and promotes metastases in triple-negative breast cancer. Oncogene. 2016;36(15):2116–2130. doi:10.1038/onc.2016.369.

11. Dong Y, Kang H, Liu H, Wang J, Guo Q, Song C, et al. Myoferlin, a Membrane Protein with Emerging Oncogenic Roles. BioMed Research International. 2019;2019:1–9. doi:10.1155/2019/7365913.

12. Huang Y, Han Y, Guo R, Liu H, Li X, Jia L, et al. Long non-coding RNA FER1L4 promotes osteogenic differentiation of human periodontal ligament stromal cells via miR-874-3p and vascular endothelial growth factor A. Stem Cell Research & Therapy. 2020;11(1). doi:10.1186/s13287-019-1519-z.

13. Ouyang L, Yang M, Wang X, Fan J, Liu X, Zhang Y, et al. Long non-coding RNA FER1L4 inhibits cell proliferation and promotes cell apoptosis via the PTEN/AKT/p53 signaling pathway in lung cancer. Oncology Reports. 2020;45(1):359–367. doi:10.3892/or.2020.7861.

14. Gao X, Wang N, Wu S, Cui H, An X, Yang Y. Long non-coding RNA FER1L4 inhibits cell proliferation and metastasis through regulation of the PI3K/AKT signaling pathway in lung cancer cells. Molecular Medicine Reports. 2019;doi:10.3892/mmr.2019.10219.

15. Fei D, Zhang X, Liu J, Tan L, Xing J, Zhao D, et al. Long Noncoding RNA FER1L4 Suppresses Tumorigenesis by Regulating the Expression of PTEN Targeting miR-18a-5p in Osteosarcoma. Cellular Physiology and Biochemistry. 2018;51(3):1364–1375. doi:10.1159/000495554.

16. You Z, Ge A, Pang D, Zhao Y, Xu S. Long noncoding RNA FER1L4 acts as an oncogenic driver in human pan-cancer. Journal of Cellular Physiology. 2019;235(2):1795–1807. doi:10.1002/jcp.29098.

17. Liu S, Zou B, Tian T, Luo X, Mao B, Zhang X, et al. Overexpression of the lncRNA FER1L4 inhibits paclitaxel tolerance of ovarian cancer cells via the regulation of the MAPK signaling pathway. Journal of Cellular Biochemistry. 2018;120(5):7581–7589. doi:10.1002/jcb.28032.

18. Huo W, Qi F, Wang K. Long non-coding RNA FER1L4 inhibits prostate cancer progression via sponging miR-92a-3p and upregulation of FBXW7. Cancer Cell International. 2020;20(1). doi:10.1186/s12935-020-1143-0.

19. Kalyani RU, Perinbam K, Jeyanthi P, Al-Dhabi NA, Arasu MV, Esmail GA, et al. Fer1L5, a Dysferlin Homologue Present in Vesicles and Involved in C2C12 Myoblast Fusion and Membrane Repair. Biology. 2020;9(11):386. doi:10.3390/biology9110386.

20. Posey AD, Pytel P, Gardikiotes K, Demonbreun AR, Rainey M, George M, et al. Endocytic Recycling Proteins EHD1 and EHD2 Interact with Fer-1-like-5 (Fer1L5) and Mediate Myoblast Fusion. Journal of Biological Chemistry. 2011;286(9):7379–7388. doi:10.1074/jbc.m110.157222.

21. Ramachandran UK. Characterization of Fer1L5, a novel dysferlin myoferlin related protein. Durham University Dissertation. 2009;.

22. Bonventre JA, Holman C, Manchanda A, Codding SJ, Chau T, Huegel J, et al. Fer1L6 is essential for the development of vertebrate muscle tissue in zebrafish. Molecular Biology of the Cell. 2019;30(3):293–301. doi:10.1091/mbc.e18-06-0401.

23. Wolk C. Initial *in vitro* and *in vivo* Characterization of the Membrane Trafficking Protein Fer1L6. Oregon State University Dissertation. 2017;.

24. Redpath GMI, Sophocleous RA, Turnbull L, Whitchurch CB, Cooper ST. Ferlins Show Tissue-Specific Expression and Segregate as Plasma Membrane/Late Endosomal or Trans-Golgi/Recycling Ferlins. Traffic. 2016;17(3):245–266. doi:10.1111/tra.12370.

25. Sutton RB, Davletov BA, Berghuis AM, Sudhof TC, Sprang SR. Structure of the first C2 domain of synaptotagmin 1: A novel Ca^2+^/phospholipid-binding fold. Cell. 1995;80(6):929–938. doi:10.1016/0092-8674(95)90296-1.

26. Corbalan-Garcia S, Gómez-Fernández JC. Signaling through C2 domains: More than one lipid target. Biochimica et Biophysica Acta (BBA) - Biomembranes. 2014;1838(6):1536–1547. doi:10.1016/j.bbamem.2014.01.008.

27. Shao X, Davletov BA, Sutton RB, Südhof TC, Rizo J. Bipartite Ca^2+^-Binding Motif in C2 Domains of Synaptotagmin and Protein Kinase C. Science. 1996;273(5272):248–251. doi:10.1126/science.273.5272.248.

28. Ubach J, Zhang X, Shao X, Südhof TC, Rizo J. Ca^2+^ binding to synaptotagmin: how many calcium ions bind to the tip of a C2-domain? The EMBO Journal. 1998;17(14):3921–3930. doi:10.1093/emboj/17.14.3921.

29. Davis AF, Bai J, Fasshauer D, Wolowick MJ, Lewis JL, Chapman ER. Kinetics of Synaptotagmin Responses to Ca^2+^ and Assembly with the Core SNARE Complex onto Membranes. Neuron. 1999;24(2):363–376. doi:10.1016/s0896-6273(00)80850-8.

30. Bashir R, Britton S, Strachan T, Keers S, Vafiadaki E, Lako M, et al. A gene related to *Caenorhabditis elegans* spermatogenesis factor *fer-1* is mutated in Limb-Girdle Muscular Dystrophy-type 2B. Nature Genetics. 1998;20(1):37–42. doi:10.1038/1689.

31. Britton S, Freeman T, Vafiadaki E, Keers S, Harrison R, Bushby K, et al. The Third Human FER-1-like Protein Is Highly Similar to Dysferlin. Genomics. 2000;68(3):313–321. doi:10.1006/geno.2000.6290.

32. Therrien C, Dodig D, Karpati G, Sinnreich M. Mutation impact on dysferlin inferred from database analysis and computer-based structural predictions. Journal of the Neurological Sciences. 2006;250(1):71–78. doi:https://doi.org/10.1016/j.jns.2006.07.004.

33. Therrien C, Fulvio SD, Pickles S, Sinnreich M. Characterization of Lipid Binding Specificities of Dysferlin C2 Domains Reveals Novel Interactions with Phosphoinositides. Biochemistry. 2009;48(11):2377–2384. doi:10.1021/bi802242r.

34. Abdullah N, Padmanarayana M, Marty NJ, Johnson CP. Quantitation of the Calcium and Membrane Binding Properties of the C2 Domains of Dysferlin. Biophysical Journal. 2014;106(2):382–389. doi:10.1016/j.bpj.2013.11.4492.

35. Llanga T, Nagy N, Conatser L, Dial C, Sutton RB, Hirsch ML. Structure-Based Designed Nano-Dysferlin Significantly Improves Dysferlinopathy in BLA/J Mice. Molecular Therapy. 2017;25(9):2150–2162. doi:10.1016/j.ymthe.2017.05.013.

36. Jumper J, Evans R, Pritzel A, Green T, Figurnov M, Ronneberger O, et al. Highly accurate protein structure prediction with AlphaFold. Nature. 2021;596(7873):583–589. doi:10.1038/s41586-021-03819-2.

37. Baek M, DiMaio F, Anishchenko I, Dauparas J, Ovchinnikov S, Lee GR, et al. Accurate prediction of protein structures and interactions using a three-track neural network. Science. 2021;373(6557):871–876. doi:10.1126/science.abj8754.

38. Anfinsen CB. Principles that Govern the Folding of Protein Chains. Science. 1973;181(4096):223–230. doi:10.1126/science.181.4096.223.

39. Davletov BA, Südhof TC. A single C2 domain from synaptotagmin I is sufficient for high affinity Ca^2+^/phospholipid binding. J Biol Chem. 1993;268(35):26386–26390.

40. Eswar N, Webb B, Marti-Renom MA, Madhusudhan MS, Eramian D, yi Shen M, et al. Comparative Protein Structure Modeling Using Modeller. Current Protocols in Bioinformatics. 2006;15(1). doi:10.1002/0471250953.bi0506s15.

41. Nalefski EA, Falke JJ. The C2 domain calcium-binding motif: Structural and functional diversity. Protein Science. 1996;5(12):2375–2390. doi:10.1002/pro.5560051201.

42. Lee J, Cheng X, Swails JM, Yeom MS, Eastman PK, Lemkul JA, et al. CHARMM-GUI Input Generator for NAMD, GROMACS, AMBER, OpenMM, and CHARMM/OpenMM Simulations Using the CHARMM36 Additive Force Field. Journal of Chemical Theory and Computation. 2015;12(1):405–413. doi:10.1021/acs.jctc.5b00935.

43. Phillips JC, Hardy DJ, Maia JDC, Stone JE, Ribeiro JV, Bernardi RC, et al. Scalable molecular dynamics on CPU and GPU architectures with NAMD. The Journal of Chemical Physics. 2020;153(4):044130. doi:10.1063/5.0014475.

44. Waterhouse A, Bertoni M, Bienert S, Studer G, Tauriello G, Gumienny R, et al. SWISS-MODEL: homology modelling of protein structures and complexes. Nucleic Acids Research. 2018;46(W1):W296–W303. doi:10.1093/nar/gky427.

45. Pei J, Kim BH, Grishin NV. PROMALS3D: a tool for multiple protein sequence and structure alignments. Nucleic Acids Research. 2008;36(7):2295–2300. doi:10.1093/nar/gkn072.

46. Robert X, Gouet P. Deciphering key features in protein structures with the new ENDscript server. Nucleic Acids Research. 2014;42(W1):W320–W324. doi:10.1093/nar/gku316.

47. Hirsch ML, Li C, Bellon I, Yin C, Chavala S, Pryadkina M, et al. Oversized AAV Transductifon Is Mediated via a DNA-PKcs-independent, Rad51C-dependent Repair Pathway. Molecular Therapy. 2013;21(12):2205–2216. doi:10.1038/mt.2013.184.

48. Micsonai A, Wien F, Kernya L, Lee YH, Goto Y, Réfrégiers M, et al. Accurate secondary structure prediction and fold recognition for circular dichroism spectroscopy. Proceedings of the National Academy of Sciences. 2015;112(24):E3095–E3103. doi:10.1073/pnas.1500851112.

49. Micsonai A, Wien F, Bulyáki É, Kun J, Moussong É, Lee YH, et al. BeStSel: a web server for accurate protein secondary structure prediction and fold recognition from the circular dichroism spectra. Nucleic Acids Research. 2018;46(W1):W315–W322. doi:10.1093/nar/gky497.

50. Drew ED, Janes RW. PDBMD2CD: providing predicted protein circular dichroism spectra from multiple molecular dynamics-generated protein structures. Nucleic Acids Research. 2020;48(W1):W17–W24. doi:10.1093/nar/gkaa296.

51. Lek A, Lek M, North KN, Cooper ST. Phylogenetic analysis of ferlin genes reveals ancient eukaryotic origins. BMC Evolutionary Biology. 2010;10(1):231. doi:10.1186/1471-2148-10-231.

52. Fuson K, Rice A, Mahling R, Snow A, Nayak K, Shanbhogue P, et al. Alternate Splicing of Dysferlin C2A Confers Ca^2+^-Dependent and Ca^2+^-Independent Binding for Membrane Repair. Structure. 2014;22(1):104–115. doi:10.1016/j.str.2013.10.001.

53. Harsini FM, Bui AA, Rice AM, Chebrolu S, Fuson KL, Turtoi A, et al. Structural Basis for the Distinct Membrane Binding Activity of the Homologous C2A Domains of Myoferlin and Dysferlin. Journal of Molecular Biology. 2019;431(11):2112–2126. doi:10.1016/j.jmb.2019.04.006.

54. Wang Y, Tadayon R, Santamaria L, Mercier P, Forristal CJ, Shaw GS. Calcium binds and rigidifies the dysferlin C2A domain in a tightly coupled manner. Biochemical Journal. 2021;478(1):197–215. doi:10.1042/bcj20200773.

55. Helfmann S, Neumann P, Tittmann K, Moser T, Ficner R, Reisinger E. The Crystal Structure of the C2A Domain of Otoferlin Reveals an Unconventional Top Loop Region. Journal of Molecular Biology. 2011;406(3):479–490. doi:10.1016/j.jmb.2010.12.031.

56. Nagashima T, Hayashi F, and SY. Solution structure of the first C2 domain of human myoferlin. PDB Entry - 2DMH. 2006;doi:10.2210/pdb2dmh/pdb.

57. Otto SC, Reardon PN, Kumar TM, Kuykendall CJ, Johnson CP. Calcium Sensitive Allostery Regulates the PI(4,5)P2 Binding Site of the Dysferlin C2A Domain. bioRxiv. 2021;doi:10.1101/2021.02.10.430549.

58. Blom N, Gammeltoft S, Brunak S. Sequence and structure-based prediction of eukaryotic protein phosphorylation sites. Journal of Molecular Biology. 1999;294(5):1351–1362. doi:10.1006/jmbi.1999.3310.

59. Pramono ZAD, Tan CL, Seah IAL, See JSL, Kam SY, Lai PS, et al. Identification and characterisation of human dysferlin transcript variants: implications for dysferlin mutational screening and isoforms. Human Genetics. 2009;125(4):413–420. doi:10.1007/s00439-009-0632-y.

60. Dessen A, Tang J, Schmidt H, Stahl M, Clark JD, Seehra J, et al. Crystal Structure of Human Cytosolic Phospholipase A2 Reveals a Novel Topology and Catalytic Mechanism. Cell. 1999;97(3):349–360. doi:10.1016/s0092-8674(00)80744-8.

61. Harsini FM, Chebrolu S, Fuson KL, White MA, Rice AM, Sutton RB. FerA is a Membrane-Associating Four-Helix Bundle Domain in the Ferlin Family of Membrane-Fusion Proteins. Scientific Reports. 2018;8(1). doi:10.1038/s41598-018-29184-1.

62. Sula A, Cole AR, Yeats C, Orengo C, Keep NH. Crystal structures of the human Dysferlin inner DysF domain. BMC Structural Biology. 2014;14(1). doi:10.1186/1472-6807-14-3.

63. Patel P, Harris R, Geddes SM, Strehle EM, Watson JD, Bashir R, et al. Solution Structure of the Inner DysF Domain of Myoferlin and Implications for Limb Girdle Muscular Dystrophy Type 2B. Journal of Molecular Biology. 2008;379(5):981–990. doi:10.1016/j.jmb.2008.04.046.

64. Yan M, Rachubinski DA, Joshi S, Rachubinski RA, Subramani S. Dysferlin Domain-containing Proteins, Pex30p and Pex31p, Localized to Two Compartments, Control the Number and Size of Oleate-induced Peroxisomes in Pichia pastoris. Molecular Biology of the Cell. 2008;19(3):885–898. doi:10.1091/mbc.e07-10-1042.

65. Vizeacoumar FJ, Torres-Guzman JC, Bouard D, Aitchison JD, Rachubinski RA. Pex30p, Pex31p, and Pex32p Form a Family of Peroxisomal Integral Membrane Proteins Regulating Peroxisome Size and Number in *Saccharomyces cerevisiae*. Molecular Biology of the Cell. 2004;15(2):665–677. doi:10.1091/mbc.e03-09-0681.

66. Okumura Y, Nakamura TS, Tanaka T, Inoue I, Suda Y, Takahashi T, et al. The Dysferlin Domain-Only Protein, Spo73, Is Required for Prospore Membrane Extension in *Saccharomyces cerevisiae*. mSphere. 2016;1(1). doi:10.1128/msphere.00038-15.

67. Wetzel L, Blanchard S, Rama S, Beier V, Kaufmann A, Wollert T. TECPR1 promotes aggrephagy by direct recruitment of LC3C autophagosomes to lysosomes. Nature Communications. 2020;11(1). doi:10.1038/s41467-020-16689-5.

68. Espinoza-Fonseca LM. Pathogenic mutation R959W alters recognition dynamics of dysferlin inner DysF domain. Molecular BioSystems. 2016;12(3):973–981. doi:10.1039/c5mb00772k.

69. Lin YF, Cheng CW, Shih CS, Hwang JK, Yu CS, Lu CH. MIB: Metal Ion-Binding Site Prediction and Docking Server. Journal of Chemical Information and Modeling. 2016;56(12):2287–2291. doi:10.1021/acs.jcim.6b00407.

70. Redpath GMI, Woolger N, Piper AK, Lemckert FA, Lek A, Greer PA, et al. Calpain cleavage within dysferlin exon 40a releases a synaptotagmin-like module for membrane repair. Molecular Biology of the Cell. 2014;25(19):3037–3048. doi:10.1091/mbc.e14-04-0947.

71. Fuson KL, Montes M, Robert JJ, Sutton RB. Structure of human synaptotagmin 1 C2AB in the absence of Ca^2+^ reveals a novel domain association. Biochemistry. 2007;46(45):13041–13048.

72. Bycroft M, Grünert S, Murzin AG, Proctor M, St Johnston D. NMR solution structure of a dsRNA binding domain from *Drosophila* staufen protein reveals homology to the N-terminal domain of ribosomal protein S5. EMBO J. 1995;14(14):3563–3571.

73. Wilson RC, Tambe A, Kidwell MA, Noland CL, Schneider CP, Doudna JA. Dicer-TRBP Complex Formation Ensures Accurate Mammalian MicroRNA Biogenesis. Molecular Cell. 2015;57(3):397–407. doi:10.1016/j.molcel.2014.11.030.

74. Moller S, Croning MDR, Apweiler R. Evaluation of methods for the prediction of membrane spanning regions. Bioinformatics. 2001;17(7):646–653. doi:10.1093/bioinformatics/17.7.646.

75. Middel V, Zhou L, Takamiya M, Beil T, Shahid M, Roostalu U, et al. Dysferlin-mediated phosphatidylserine sorting engages macrophages in sarcolemma repair. Nature Communications. 2016;7(1). doi:10.1038/ncomms12875.

76. Varga R. OTOF mutations revealed by genetic analysis of hearing loss families including a potential temperature sensitive auditory neuropathy allele. Journal of Medical Genetics. 2005;43(7):576–581. doi:10.1136/jmg.2005.038612.

77. Cacciottolo M, Numitone G, Aurino S, Caserta IR, Fanin M, Politano L, et al. Muscular dystrophy with marked Dysferlin deficiency is consistently caused by primary dysferlin gene mutations. European Journal of Human Genetics. 2011;19(9):974–980. doi:10.1038/ejhg.2011.70.

78. Argon Y, Ward S. *Caenorhabditis elegans* fertilization-defective mutants with abnormal sperm. Genetics. 1980;96(2):413–433.

79. Washington NL, Ward S. FER-1 regulates Ca^2+^-mediated membrane fusion during *C. elegans* spermatogenesis. J Cell Sci. 2006;119(Pt 12):2552–2562.

80. Krajacic P, Hermanowski J, Lozynska O, Khurana TS, Lamitina T. *C. elegans* dysferlin homolog *fer-1* is expressed in muscle, and *fer-1* mutations initiate altered gene expression of muscle enriched genes. Physiol Genomics. 2009;40(1):8–14.

81. Lek A, Evesson FJ, Sutton RB, North KN, Cooper ST. Ferlins: Regulators of Vesicle Fusion for Auditory Neurotransmission, Receptor Trafficking and Membrane Repair. Traffic. 2011;13(2):185–194. doi:10.1111/j.1600-0854.2011.01267.x.

82. Johnson CP, Chapman ER. Otoferlin is a calcium sensor that directly regulates SNARE-mediated membrane fusion. Journal of Cell Biology. 2010;191(1):187–197. doi:10.1083/jcb.201002089.

83. Perisic O, Fong S, Lynch DE, Bycroft M, Williams RL. Crystal Structure of a Calcium-Phospholipid Binding Domain from Cytosolic Phospholipase A2. Journal of Biological Chemistry. 1998;273(3):1596–1604. doi:10.1074/jbc.273.3.1596.

